# Modulation of feature attention by reward prediction error explains value learning behavior

**DOI:** 10.64898/2026.04.10.717847

**Authors:** Mingze L. Leukos (李铭泽), Albert Liang, Grace W. Lindsay

**Author notes:** Corresponding Author: Grace W. Lindsay. **Conflict of Interest:** The authors declare no competing interests. **Author Contributions:** ML contributed to designing the research, performed the research, analyzed data and wrote the first draft of the paper. AL contributed to data analysis. GWL designed the research and edited the paper.

## Abstract

Adaptive behavior requires learning the value of environmental features while selectively attending to those most likely to yield reward. Reward prediction errors (RPEs) drive value learning and learned values guide attention, yet the computational function linking RPEs to attentional modulation remains unspecified. Here, we developed a reinforcement learning model with a perceptual front-end to investigate how value and RPE signals modulate attentional gain during learning. We compared five candidate RPE-attention transfer functions, each combined with either single- or multi-focus attention, against behavioral data from two adult male rhesus macaques performing a color-value learning task with shifting reward contingencies. Monkeys exhibited rapid initial learning followed by sub-optimal asymptotic accuracy. Overall, single-focus architectures consistently outperformed multi-focus counterparts on matching monkey errors, indicating that macaques collapse the value distribution into a winner-take-all attentional focus. Furthermore, the “Switch” model, in which attention targets the highest-valued feature but transiently inverts following negative RPEs, produced the fastest exploration dynamics following target switches and, together with the Absolute Value model, yielded decision confidence trajectories that positively correlated with empirical reaction times. In support of this, single-neuron correlation analyses revealed that 27 − 42% of neurons in prefrontal cortex, frontal eye fields, and lateral intraparietal area encoded previous-trial RPE at the time of next trial onset. In total, we conclude that capacity constrained attention that inverts its focus after negative RPE best explains value learning dynamics. These results provide a normative account for why biological learners sacrifice asymptotic precision for rapid adaptation in volatile environments.

**Significance Statement:** Learning which features of the environment predict reward requires both reinforcement learning and selective attention, yet how these processes interact algorithmically remains unknown. We developed a computational model that specifies how reward prediction errors dynamically adjust the gain of feature-based attention during value learning. Testing competing hypotheses against macaque behavioral data, we show that a “Switch” mechanism, in which negative prediction errors transiently invert attentional focus away from the highest-valued feature, best captures primate learning dynamics. This architecture reveals a principled trade-off: the brain sacrifices asymptotic accuracy for rapid detection of environmental change, using error-triggered attentional inversion as a directed exploration strategy. These findings bridge reinforcement learning theory and attention research by identifying the transfer function linking prediction errors to sensory gain modulation.

## 1 Introduction

To behave adaptively, organisms must learn the values of environmental features while selectively attending to those most likely to yield reward. This establishes a closed loop between reinforcement learning (RL) and feature-based attention: while rewards drive learning by updating internal value estimates (Schultz et al., 1997; Glimcher, 2011), these value estimates guide the deployment of attention to relevant features (Lindsay, 2020); attention, in turn, constrains the state representation over which the RL system operates, filtering multi-stimuli environments to prioritize reward-relevant options (Niv et al., 2015; Radulescu et al., 2019). Past value learning thus influences current attentional selection, and current attentional selection biases the action selection that guides future learning.

The learning side of this loop is formalized through temporal difference (TD) algorithms, which update value estimates based on Reward Prediction Errors (RPEs): the discrepancy between expected and received rewards (Sutton et al., 1998). These values subsequently bias action selection: learned values bias saccades toward stimuli previously associated with higher reward, even when such stimuli are no longer task-relevant (Anderson and Yantis, 2012; Hikosaka et al., 2006), a phenomenon mediated by interactions between dopaminergic reward circuits and frontoparietal attention networks (Anderson et al., 2011; Anderson, 2017). Furthermore, overt visual attention patterns predict value-driven choice behavior, demonstrating that attention also dynamically shapes the computation underlying value-based decisions rather than merely reflecting learned value (Perkins and Rich, 2024).

Mechanistically, feature-based attention provides selective enhancement of neural responses to particular stimulus attributes by modulating the gain of sensory neurons tuned to attended features (Treue and Trujillo, 1999; Maunsell and Treue, 2006; Martinez-Trujillo and Treue, 2004). This top-down modulation is controlled by frontoparietal and prefrontal networks, with the frontal eye fields (FEF) and lateral intraparietal area (LIP) maintaining spatial priority maps that direct attention to relevant locations (Buschman and Miller, 2007) and ventral pre-arcuate region of prefrontal cortex serving as a source of feature-based attentional signals that are relayed to FEF to guide selection (Bichot et al., 2015).

While it is established that value modulates attention and attention shapes learning, the specific transfer function mapping value and RPE onto attentional gain remains unknown. Emerging evidence suggests that RPE may directly regulate attentional allocation. Behaviorally, increases in prediction error broaden attentional allocation to previously irrelevant cues (Torrents-Rodas et al., 2021). In multidimensional learning environments, the absence of expected reward predicts trial-by-trial shifts in attentional focus across stimulus dimensions (Leong et al., 2017). At the neural level, feature-specific prediction error and surprise signals have been identified in macaque fronto-striatal circuits, indicating that the brain computes prediction errors in a manner that could selectively target feature-based attention (Oemisch et al., 2019). Together, these findings point to RPE as a candidate signal for dynamically regulating attention strength, but the precise mathematical relationship governing this modulation remains unspecified.

The omission of the precise mathematical relationship between RL variables and attention is particularly consequential because standard RL models often assume unbiased sensory access, failing to account for how attentional bottlenecks impact and are impacted by the learning process itself (Radulescu et al., 2019; Eckstein et al., 2021), especially in volatile environments where the learner must balance exploitation of currently high-value features against exploration in reward contingencies.

Here, we developed a perceptual reinforcement learning model where top-down feature-selective attention is allocated according to an internal value function and modulated by RPE, allowing us to test competing hypotheses regarding the RPE-value-attention relationship. We compared these architectures against behavioral data from macaques performing a color-value learning task (Jahn et al., 2024) and analyzed simultaneously recorded neural activity to examine the nature of RPE signals in frontal and parietal cortex. We found that the specific learning trajectories observed in primates, characterized by rapid initial acquisition followed by sub-optimal asymptotic performance, are best explained by a single-focus “Switch” mechanism. In this architecture, attention is focused on the highest-valued feature but is transiently inverted following negative prediction errors. This model suggests a normative account of value-based attention: the brain prioritizes the rapid extraction of high-value signals over precise probabilistic representation, accepting a ceiling on asymptotic accuracy to ensure speed of adaptation in volatile environments.

## 2 Materials and Methods

### 2.1 Color-Value Learning Task

Jahn et al. (2024) trained 2 monkeys on a color-value learning task and simultaneously recorded behavioral and neural data (see Figure 1). On each trial, three colored stimuli were drawn uniformly from a color wheel (100 evenly spaced colors) and randomly appeared at three of four fixed locations, while the monkey is fixated at center. The monkeys were trained to select one of the three stimuli by making a saccade to its location. They then received a juice reward based on how far the chosen color *c* was from an unknown target color *c*_target_ (i.e. the color with the highest value in the true color-value distribution). Specifically, reward decreased with the angular distance between the chosen color *c*_chosen_ and the hidden target color *c*_target_ on the color wheel according to:

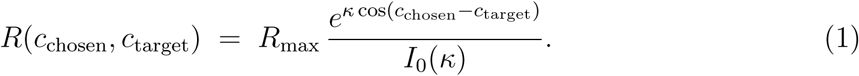

**Figure 1:**
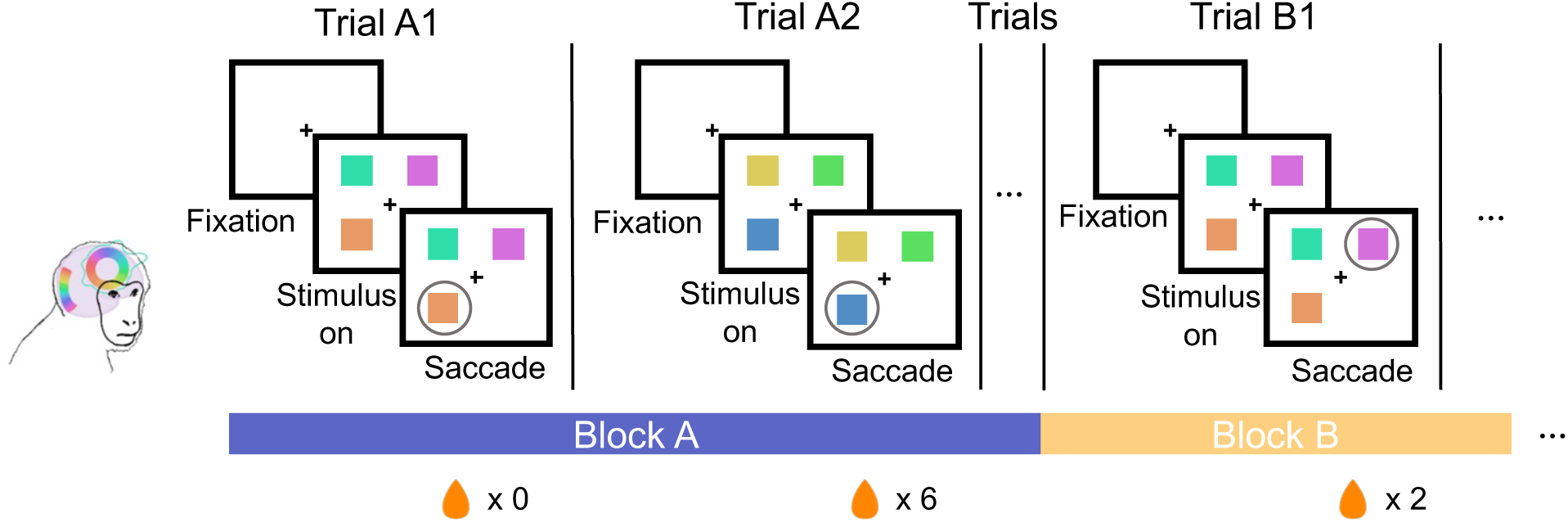
Color-value learning task from (Jahn et al., 2024) On each trial, three colored stimuli were shown while the monkey fixated centrally. The monkey chose a stimulus via saccade and received a juice reward based on the stimulus color’s proximity to the target. The target color remained fixed within a block, but changed without notice after 80-200 trials. The bottom timeline illustrates two consecutive blocks (Block A, Block B) with their respective target colors.

where *c*_chosen_ is the chosen color, *c*_target_ is the target color, *κ* is the concentration parameter, and *I*_0_(*κ*) is the modified Bessel function of the first kind of order zero.

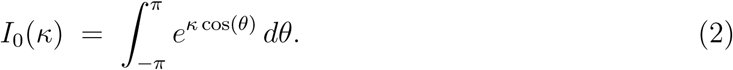

The true value distribution of the colors was thus not known to the monkey: the monkey had to estimate the colors’ values through reward feedback.

Critically, the target color *c*_target_ changes without notice roughly every 80 - 200 trials. The change was triggered when the monkey chose the best available color on at least 80% (Monkey B) or 85% (Monkey S) of 30 consecutive trials (Jahn et al., 2024). We refer to the set of trials in between a target change (wherein the target is constant) as a block. The maximum reward *R*_max_ was fixed at 6 drops for Monkey S but increased by 1 drop per completed block for Monkey B (starting at 12 drops), creating an escalating reward schedule for Monkey B only (Jahn et al., 2024).

For our model, we replicate the task setup exactly by using the same stimuli shown to the monkey on each trial. We therefore switch the target color *c*_target_ on the same trials as they occur for the monkeys (regardless of model performance). For reward, we use the same reward function (1). In the original experiment, reward magnitude (delivered as juice drops) was rounded to the nearest integer before delivery to the monkey (Jahn et al., 2024); in our simulations, we use the continuous (unrounded) reward function. We normalize the reward function (which is originally in units of juice) to have 1.0 as the maximum reward (varying the maximum reward to other values in the range of [0.5, 3.0] yielded similar results).

### 2.2 Model Training and Simulation

#### Value Function Learning

The model learns the value function *V* (*c*) through reinforcement learning using temporal difference (TD) learning. Following Q-learning methods in Jahn et al. (2024), the value function is represented using a set of *N* equally distant radial basis functions centered around the color wheel. (*N* = 6 as in Jahn et al. (2024)). The value function is defined over all colors *c* on the color wheel, allowing the model to estimate the expected reward for any stimulus (see Figure 2):

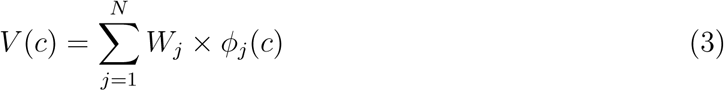

**Figure 2:**
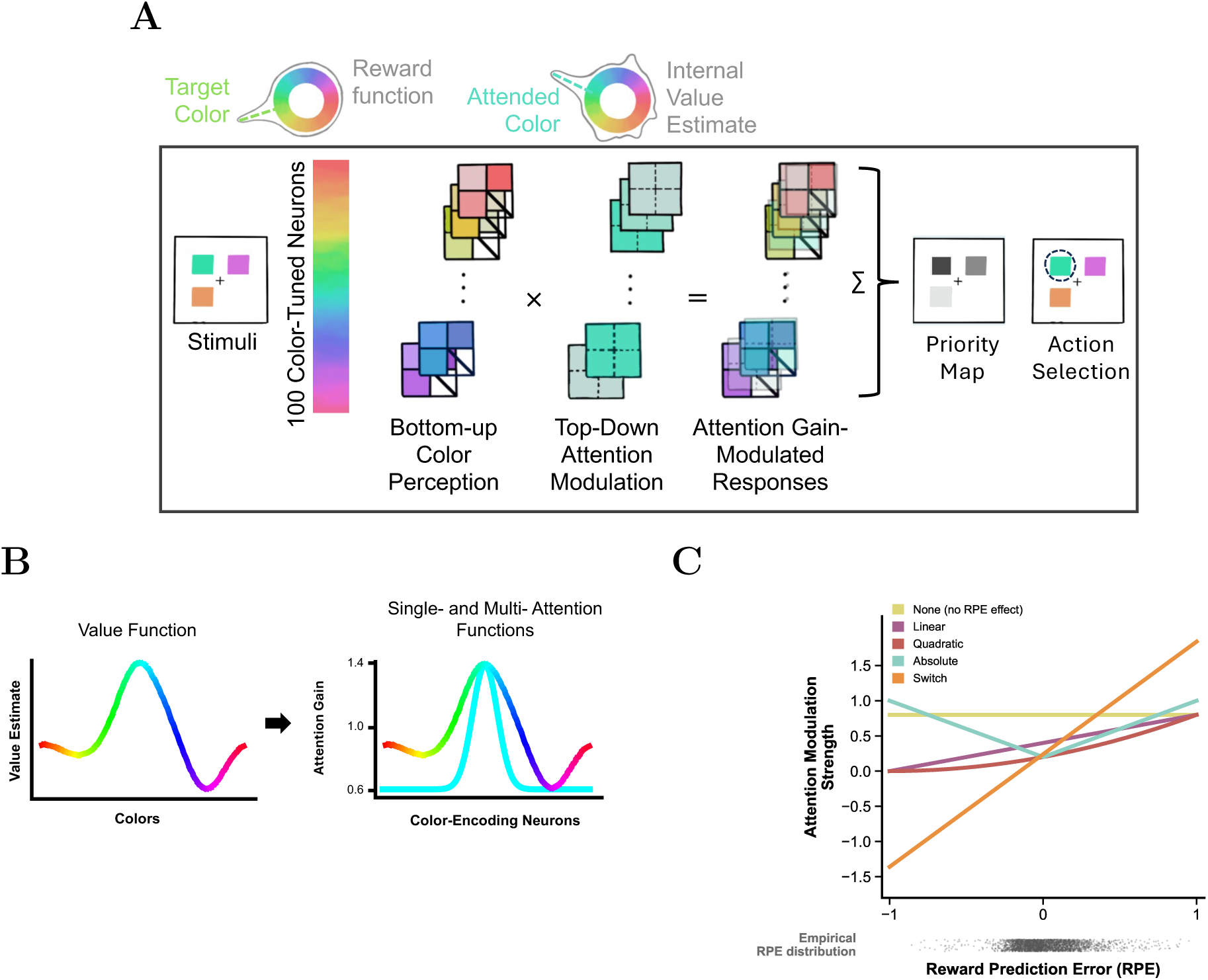
Model architecture and attention function options. **(A)** Schematic of the perceptual reinforcement learning model. The model maintains an internal value estimate over the color wheel. Bottom-up responses of color-tuned neurons are multiplicatively modulated by a top-down attentional gain signal derived from the value estimate (see panel B) and scaled by previous-trial RPE (see panel C). The gain-modulated responses are summed across neurons at each location to produce a priority map, from which an action is selected stochastically. **(B)** Conversion of the internal value function (left) into attentional gain profiles (right). In the single-focus mechanism (cyan), attention is concentrated on the color with the highest value. In the multi-focus mechanism (rainbow), attentional gain is distributed across all colors in proportion to their learned values. **(C)** RPE-attention transfer functions. Each line shows how the attention modulation strength varies as a function of previous-trial RPE for the five candidate transfer functions. The scatter strip below the x-axis shows the empirical distribution of trial-wise RPE values derived from the Q-learning model fit to monkey choice behavior.

where *ϕ_j_*(*c*) are von Mises basis functions with concentration parameter *κ* = 2.23 (as in Jahn et al. (2024)) and *W_j_* is the learned weights for basis function *j*.

In Jahn et al. (2024), several explicit biases were added to the value estimates in order to better capture the animals’ behavior. This included color preference, location bias, and previous-choice bias, as well as a reset mechanism that allowed the value function to return to uniform following large reward prediction errors. Importantly, in order to cleanly isolate the impacts of the different mechanistic hypotheses we put forth here, we do not include these additional tweaks to the value learning process.

#### Weight Update

On each trial *t*, the value function *V* (*c*) is computed for all colors *c* on the color wheel (Eq. 3). After the model selects a stimulus, weights are updated based on the reward prediction error (RPE), *δ_t_*, computed at the chosen color *c*_chosen_:

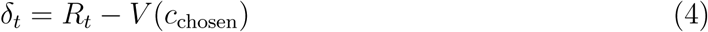

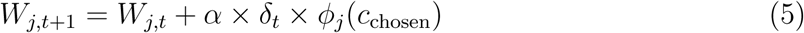

where *α* is the learning rate (set to 1.0 here unless otherwise specified), *R_t_* is the reward received, and *ϕ_j_*(*c*_chosen_) is the basis function encoding of the chosen color.

### 2.3 Perceptual Front-End

We built a perceptual front-end to enact a mechanistic model of action selection in this task. Our perceptual front-end was designed to mimic color processing in the visual system (e.g. area V4 (Bohon et al., 2016)) and serves as the target for attentional modulation (described in the following section, see Figure 2).

To simulate trials, from the data we use the exact color stimuli shown to the monkeys for each trial. Stimuli were processed through a bank of *N* = 100 color-tuned neurons, each preferring a specific color *c*_pref_ on a discretized color wheel:

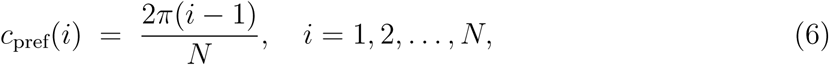

Color-tuned neuron responses were determined according to a cosine-squared color tuning function, centered on the neuron’s preferred color (as is commonly used to fit color-tuned cell responses in the visual system (Bohon et al., 2016)). Each neuron’s response *r* is calculated according to:

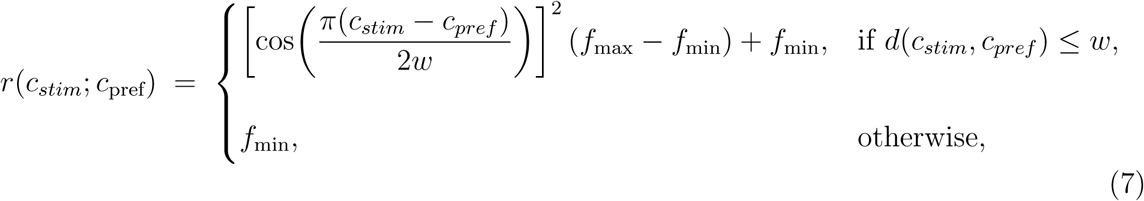

where *c*_stim_ is the color of the presented stimulus, *w* is the tuning width, *f*_max_ and *f*_min_ are the maximum and minimum firing rates, and *d*(*c*_stim_*, c*_pref_) denotes the angular distance between *c*_stim_ and *c*_pref_.

Results shown in the present report are calculated using the following tuning curve parameters: *w* = *π* radians, *f*_max_= 1.0, and *f*_min_ = 0.0. This configuration ensured that neurons showed maximal response (1.0) to their preferred color *c*_pref_, with responses decreasing according to the cosine-squared function as the angular distance from the preferred color increased, reaching zero response for colors *π* radians away. The wide tuning width (*π* radians) meant that each neuron responded to many colors, which is consistent with broad color tuning observed in the visual cortex (Sanada et al., 2016).

On each trial, model neural responses were computed independently for each of the three colored stimuli presented. Stimulus responses were organized on a 2×2 grid corresponding to the four possible stimulus locations, resulting in a 2×2×100 tensor. Absent locations were coded as “non-stimulus” and led to value of 0 in the neurons at the corresponding location.

We also compare to a baseline model where there is no perceptual frontend. Instead of computing color-tuned neuron responses and applying attentional modulation (see Value-Guided Attentional Modulation), this model operated directly on the learned value function. For each stimulus, the model retrieved the internal value estimate for the corresponding color. Action selection then proceeded by applying softmax normalization directly to these four location values (see Action Selection subsection). This baseline model thus represents a non-mechanistic simplified decision-making process that relies purely on learned internal value estimate of color features, without the intermediate step of bottom-up color perception and top-down feature-selective attentional modulation. This comparison allows us to isolate the added explanatory power of modeling the attention and sensory mechanisms that drive choice.

### 2.4 Value-Guided Attentional Modulation

The bottom-up perceptual response, processed in the perceptual front-end, was modulated by a top-down attentional signal. Consistent with experimental findings, attention was modeled as multiplicative gain modulation of the bottom-up response (Martinez-Trujillo and Treue, 2005)

For each color-tuned neuron, we calculated an attentional gain factor (*a_i_*). These attentional gain factors determined how strongly each neuron’s response was amplified or attenuated. These gain factors were calculated as a function of internal value estimates and could be modulated by previous-trial RPE (see section below).

Specifically, attended color(s) were determined by the value function via two possible ways. In the single-focus attention mechanism, a single attended color was defined as the color with the current highest internal value estimate. Attentional modulation for each neuron was then computed using a cosine-squared function (7) centered at the attended color. Psychophysical studies have shown that feature-based attention to a specific color enhances sensitivity at the attended color while suppressing perception of other colors (Wang et al., 2015). To replicate this biological effect, we set a narrow attentional width *w* = *π/*3. *f_max_* = 1.4 and *f_min_* = 0.6 represent upper and lower bound for the attention function, respectively. Thus, neurons preferring colors similar to the attended color receive attentional enhancement (*a >* 1), and neurons preferring colors farther from the attended color will receive attentional suppression (*a <* 1) (Treue and Trujillo, 1999).

In contrast, in the the multi-focus attention mechanism, attentional gain was distributed across all color bins in direct proportion to their learned values, *V*. The attention weights were computed by normalizing the value function:

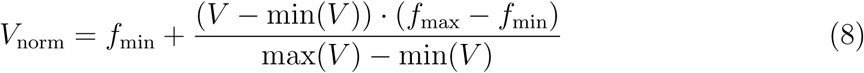

This normalization ensured that the color bin with the highest value received maximum gain (*f_max_* = 1.4), the lowest value received minimum gain (*f*_min_ = 0.6), and intermediate values received proportionally scaled gains. This enabled divided attention across multiple separate colors.

These attentional gain parameters are biologically constrained by physiological findings. Feature-based attention exerts graded multiplicative modulation on neural responses, with responses modulated by approximately 12% when attending to the preferred feature, and bounded suppression (not silencing) when the attended feature shifts from preferred to anti-preferred (Martinez-Trujillo and Treue, 2004; Mayo and Maunsell, 2016; Page and Duffy, 2008). These experimental findings support our implementation of moderate multiplicative attentional modulation that operates within bounded ranges, with both upper limits on enhancement and lower limits on suppression. Previous computational models of attention have similarly implemented bounded gain modulation to capture the constraints observed in biological attention (Lindsay and Miller, 2018). Varying the exact baseline *f*_min_ and *f*_max_ values used here leads to qualitatively similar results, but making the modulation too weak causes the differences in different attention mechanisms to be washed out by the stochasticity in action selection (see below).

The color perception map, a 2×2×100 grid map composed of all the color-tuned neurons’ bottom-up responses for all four stimulus locations, is modulated by the aforementioned attention gain(s) multiplicatively:

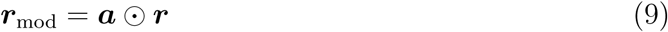

### 2.5 RPE-Guided Attention Strength

Here, inspired by past literature, we explored using the previous trial’s RPE to determine attention strength (Leong et al., 2017; Oemisch et al., 2019; Torrents-Rodas et al., 2021). This mechanism dynamically adjusted the range of the current trial’s attentional modulation based on how close the internal value estimate matched the true reward received in the previous trial, signaled by the previous trial RPE (as shown in Eq. 5, see Figure 1 panel C). The RPE-guided attention mechanism modifies the upper and lower bound of the attentional gain range, *f_max_* and *f_min_*, used in both single-focus and multi-focus attention models. We tested four mathematical relationships between previous-trial RPE (*δ_t_*) and attention strength, as follows (see Figure 2C). For each:

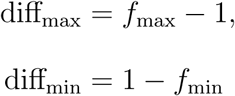

**1) Linear:**

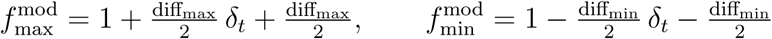

This created a linear mapping where RPE = −1 leads to minimal attention strength (approaching uniform gain), while RPE = +1 causes maximal attention strength.

**2) Quadratic:**

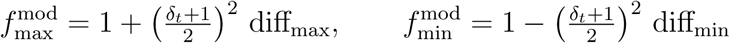

This was similar to linear but with quadratic interpolation, creating a non-linear relationship that emphasizes larger positive RPEs more strongly.

**3) Absolute Value:**

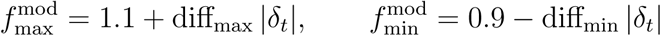

This represented unsigned RPEs, where both positive and negative prediction errors increase attention strength, reflecting the idea that any surprise should enhance focus. Offsets from 1 ensure that 0 previous-trial RPE does not lead to 0 attention modulation.

**4) Switch**

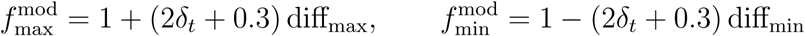

Here, negative RPEs reverse attention polarity, causing low-value stimuli to receive enhanced processing while high-value stimuli are suppressed Here again, an offset is added to ensure that 0 previous-trial RPE does not cause 0 attention modulation.

We also compare to a baseline model where attention strength is held constant and independent of RPE, representing strong attentional modulation only by value.

### 2.6 Action Selection

After applying attentional gains, the modulated responses of all color-tuned neurons at a given location are summed. This summation yields a priority map over possible actions, where locations with stimuli that match the attended features tend to result in higher summed activity and therefore receive higher priority. The priority map is normalized using total normalization, in which the value of each location is divided by the sum of all values. This ensures that the normalized values sum to 1, forming a valid probability distribution.

Action selection was implemented in either deterministic mode (selecting the location with the highest probability) or stochastic mode (locations are chosen randomly according to their probabilities). Results in the present report are all run with the stochastic mode. This design choice reflected evidence that biological decision making is inherently stochastic rather than strictly deterministic (Soltani et al., 2006)

For the baseline model without a perceptual front-end, action selection operated directly on learned stimulus values using standard softmax normalization (temperature = 0.3), following the approach and parameters optimized in (Jahn et al., 2024).

### 2.7 Simulation

In Jahn et al. (2024), there were a total of 29,874 trials. Accordingly, we simulated the same 29,874 trials using trial-wise stimulus values taken from the experimental data from Jahn et al. (2024). The model could be run in two modes: 1) closed-loop mode (where it learns from its own choices), and 2) open-loop mode (where it gets reward feedback based on the empirical choice the monkey made on each trial). Open-loop mode is used for the models in (Jahn et al., 2024). To assess the long-term stability and fidelity of our model mechanisms, results shown in the present report are all run in the closed-loop mode unless otherwise specified.

For analysis, we restrict our focus to the first 80 trials following each switch in the target color. This value was chosen because the shortest block in the monkey dataset contained 80 trials. Restricting the analysis to 80 trials aligned the model with the empirical data and enabled performance performance analysis that utilizes all blocks.

In Jahn et al. (2024), data were collected across 17 recording days from two monkeys. To mirror this setup and maintain comparability, we make the design choice such that weights, *W*, were initialized to zero at the start of each simulated day, i.e., the model makes random choices since no prior learning had occurred.

### 2.8 Learning Curve Analysis

Accuracy was computed as a binary measure on each trial, with correct choices defined as selecting the stimulus closest to the latent target color (i.e. the stimulus that nets the most reward).

To analyze learning trajectories when the target color switches, we averaged model accuracy across all blocks. The block-wise averaging is applied identically to both simulated model data and experimental behavioral data for a direct comparison between model and monkey performance.

To quantify the fit between each model’s learning trajectory and the empirical data, we computed the Mean Squared Error (MSE) between block-averaged model and monkey accuracy curves. MSE was computed separately for each of the *N* = 20 independent simulation runs (differing in random seeds), and the mean and standard error of the mean were reported across runs for each model configuration and each monkey. See Figure 3.

**Figure 3:**
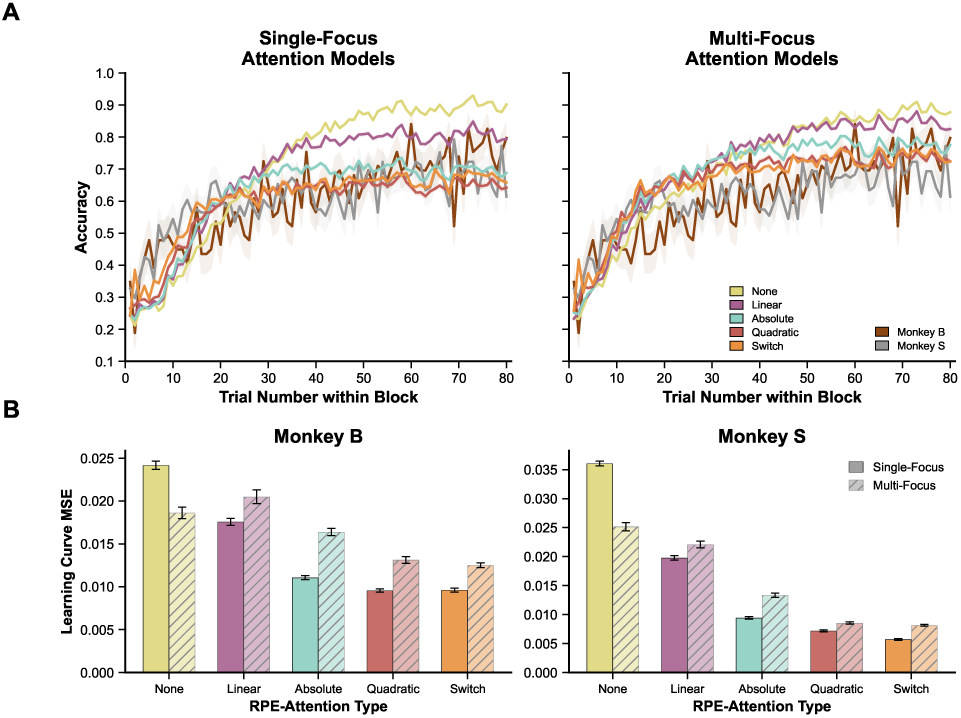
Learning curves and model fit across RPE-attention architectures. **(A)** Block-averaged accuracy (proportion of trials on which the subject selected the stimulus nearest the target color) as a function of trial number following a target color switch, for single-focus (left) and multi-focus (right) attention models. Colored lines indicate model configurations defined by RPE-attention type (None, Linear, Absolute, Quadratic, Switch); solid dark lines indicate empirical data from Monkey B (brown) and Monkey S (gray). Model curves represent the mean across *N* = 20 independent simulation runs. Monkey curves represent the mean across all blocks within each animal’s recording sessions. Shaded regions denote ±1 SEM (across runs for models; across blocks for monkeys). Chance performance is 33% (one of three stimuli). **(B)** Mean squared error (MSE) between each model’s block-averaged learning curve and the empirical monkey curve, computed separately for Monkey B (left) and Monkey S (right). Bars indicate mean MSE across *N* = 20 runs; error bars indicate ±1 SEM across runs. Solid bars, single-focus models; hatched bars, multi-focus models. Lower MSE indicates a closer fit to the empirical learning trajectory.

### 2.9 Behavioral Similarity Analysis

For additional more fine-grained comparisons between model and monkey behavior, we used four additional trial-defined behavioral similarity metrics.

**1.) Trial-wise Entropy** (abbreviated as “Entropy” in the report) The entropy metric captured the intuitive notion that trials with stimuli all of similar value should be more difficult than trials with stimuli whose values are more distributed. Therefore we defined an entropy metric that was maximized on trials where all three stimuli are equidistant from the target and minimized at trials where some stimulus is much closer or farther than another.

For each trial *t*, we compute the angular distance from each of the three stimulus colors to the target color:

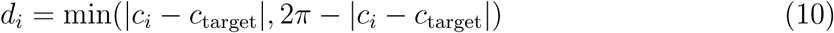

where *c_i_*is the color of stimulus *i* and *c*_target_ is the target color for that trial.

Trial entropy *H_t_*at trial t is then calculated using a softmax transformation over the inverse distances:

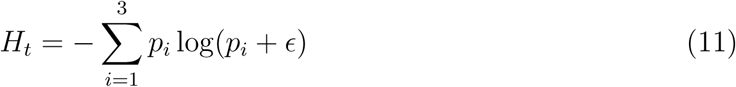

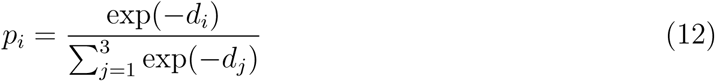

where *ɛ* = 10^−9^ prevents numerical issues with log(0). Entropy *H_t_* is maximized when all three stimuli are equidistant from the target.

**2.) Maximal Distance** The maximal distance metric captured a different aspect of task difficulty. It measured how far from the target color the farthest color stimulus is. Trials with low maximum distance values had stimuli that were clustered near the target (making it more difficult to choose the optimal one), while high maximum distance values indicate the presence of at least one stimulus far from the target(potentially making the choice easier).

For each trial *t*, we compute the angular distances *d* from all three stimulus colors to the target color.

The Maximal Distance metric for trial *t* is then defined as:

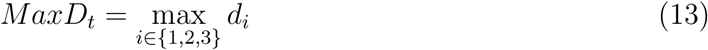

**3.) Minimal Distance** The minimal distance metric quantified how close the best available stimulus (i.e., the stimulus closest to the target color) is to the target. A high minimal distance indicated that all stimuli were far from the target color, whereas a low minimal distance means at least one is close to it (potentially making the choice easier). Similar to the Maximal Distance metric described above, we compute the minimum angular distance between the target color and any of the three stimulus options. This reflects that choosing amongst several options close to the the target color is also difficult.

**4.) Mean Distance** For the mean distance metric, we computed the mean angular distance between the target color and all three stimulus colors, under the assumption very low average and very high average trials may be equally challenging.

For each of the above four metrics, trial-wise accuracy values are binned into 20 equally-spaced bins based on the metric value, with bin means and standard errors of the mean computed for each bin. To compare model and monkey behavior, we computed the Mean Squared Error (MSE) between each model’s binned accuracy curve and the corresponding empirical monkey curve across all 20 bins. MSE was computed separately for each of the *N* = 20 independent simulation runs, and the mean and standard error of the mean were reported across runs for each model configuration and each monkey (See Figure 4).

**Figure 4:**
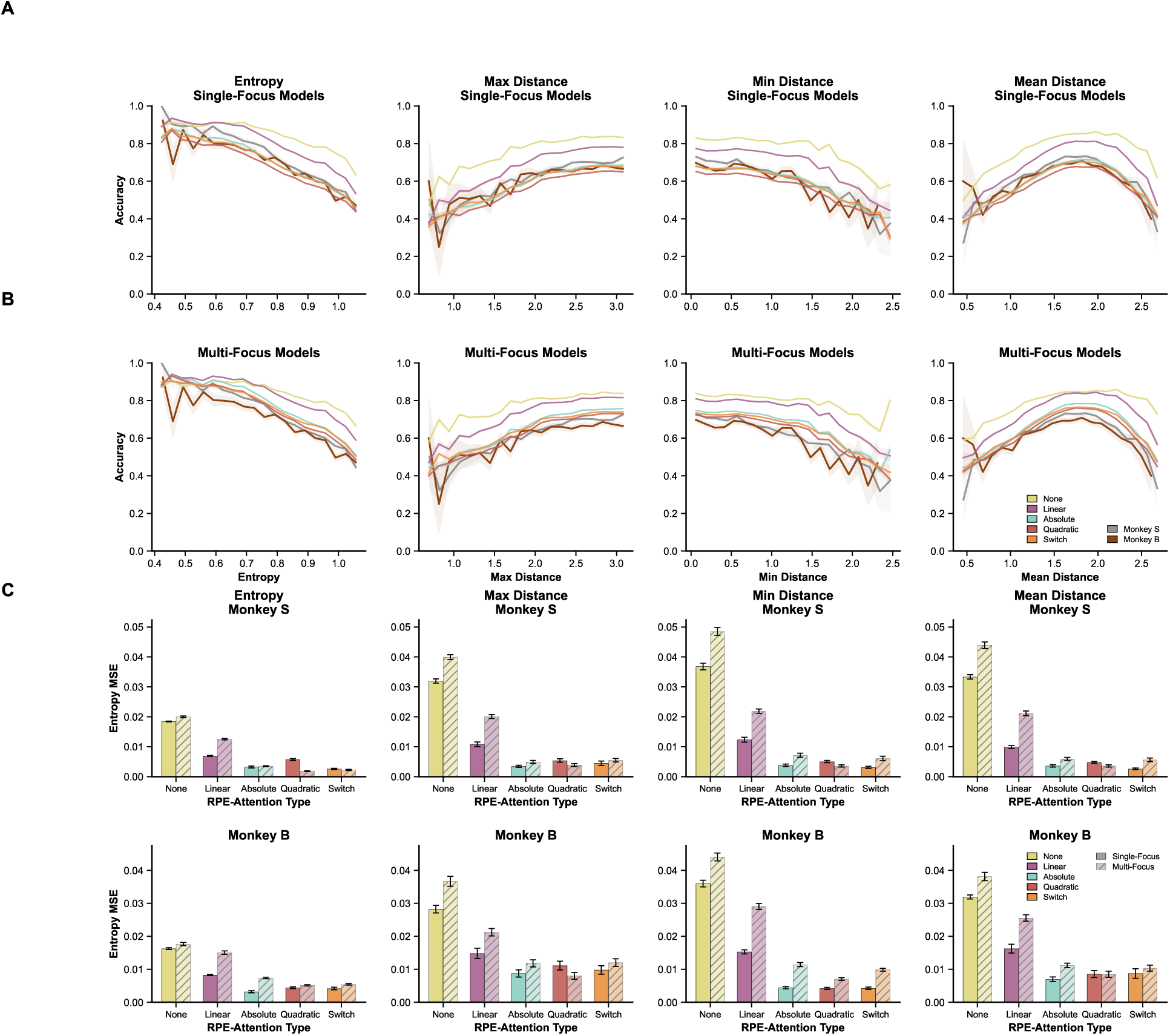
Behavioral similarity across four task-difficulty metrics. **(A)** Binned accuracy as a function of four trial-wise difficulty metrics for single-focus models: trial entropy (column 1), maximal stimulus–target angular distance (column 2), minimal stimulus–target angular distance (column 3), and mean stimulus–target angular distance column 4). For each metric, trial-wise accuracy was binned into 20 equally spaced bins. Solid colored lines indicate model accuracy averaged across *N* = 20 simulation runs; shaded regions indicate ±1 SEM. Monkey B (brown) and Monkey S (gray) empirical accuracy curves are overlaid for comparison. **(B)** Same as (A) but for multi-focus models. **(C)** MSE between binned model and monkey accuracy curves for each metric. Top row, Monkey B; bottom row, Monkey S. Bars indicate mean MSE across N=20 simulation runs; error bars indicate ±1 SEM. Solid bars, single-focus models; hatched bars, multi-focus models. Lower MSE indicates a closer match between model and monkey sensitivity to trial difficulty. Across all four metrics, single-focus architectures consistently produced lower MSE than their multi-focus counter-parts, with the recurring exception of the Quadratic condition for Monkey S.

### 2.10 Model and Animal Confidence Approximation

To explore behavior beyond just accuracy, we assessed decision confidence on each trial via two different proxy measurements. In the animals, this was simply defined as reaction time (the time between stimulus display onset and saccade initiation).

For the models, we operationalized decision confidence as the Shannon entropy of the choice probability distribution on each trial. This measure captures the degree of uncertainty in the model’s action selection: low entropy reflects a peaked distribution concentrated on a single option (high confidence), whereas high entropy reflects a diffuse distribution across alternatives (low confidence), analogous to the slower, more variable reaction times observed empirically when decision difficulty is high (Hick, 1952).

On each trial *t*, the probability assigned to each of the three available stimulus locations was extracted from the model’s action selection stage. For models incorporating the perceptual front-end, these probabilities were derived from the attention-modulated priority map following total normalization (see Action Selection). For the baseline model without a perceptual front-end, choice probabilities were taken directly from the softmax over learned stimulus values (see Figure 5).

**Figure 5:**
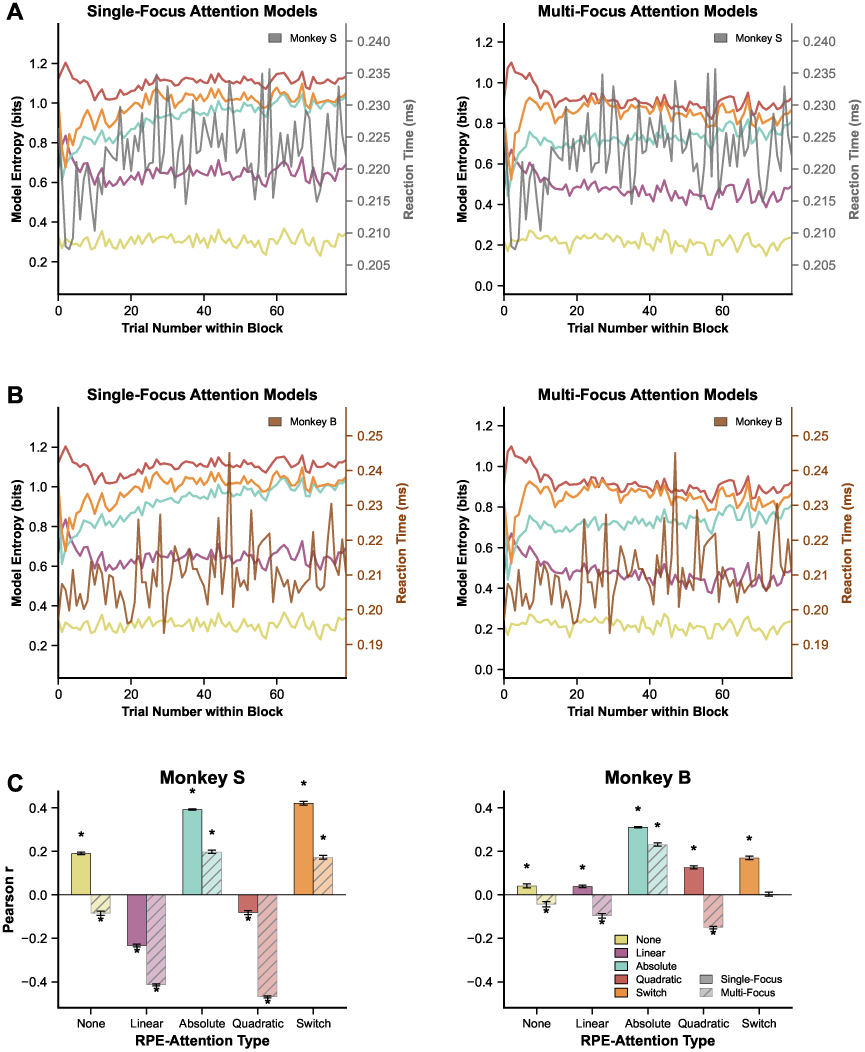
Model decision entropy correlates with empirical reaction time for Absolute and Switch RPE-attention models. **(A)** Block-averaged model decision entropy (left y-axis) and block-averaged empirical reaction time (right y-axis) as a function of trial number following a target color switch for Monkey B, shown separately for single-focus (left) and multi-focus (right) attention models. Colored lines indicate model entropy trajectories for each RPE-attention type; shaded regions denote ±1 SEM across *N* = 20 simulation runs. The brown line indicates Monkey B’s mean reaction time across blocks. Models whose entropy trajectories track the empirical RT curve (i.e., both increasing over trials within a block) produce positive Pearson correlations in panel C. **(B)** Same as (A) but for Monkey S (gray line). **(C)** Mean Pearson correlation (*r*) between block-averaged model decision entropy and block-averaged empirical reaction time, shown separately for Monkey B (left) and Monkey S (right). For each model configuration, Pearson *r* was computed per simulation run (*N* = 20) between the model’s trial-wise entropy trajectory and the monkey’s trial-wise reaction time trajectory, both block-averaged across target color switches. Bars indicate the mean Pearson *r* across runs; error bars indicate ±1 SEM. Solid bars, single-focus models; hatched bars, multi-focus models. Asterisks denote configurations for which the mean correlation differed significantly from zero (one-sample *t*-test, Benjamini-Hochberg FDR-corrected at *q <* 0.05 across the 10 model configurations tested per monkey).

Trial-wise decision entropy was then computed as:

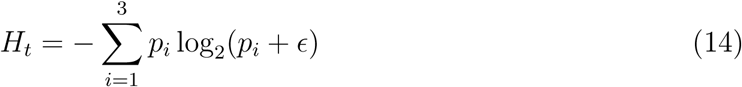

where *p_i_* is the probability assigned to stimulus location *i* and *ɛ* = 10^−12^ is a small constant added for numerical stability. *H_t_* is bounded between 0 (deterministic selection of a single option) and log_2_(3) ≈ 1.585 bits (uniform distribution across all three options).

Block-averaged entropy trajectories and their standard errors of the mean were computed across all blocks, following the same procedure applied to the empirical reaction time data. To evaluate model fit according to these different proxy measures, block-averaged entropy trajectories were compared to empirical block-averaged reaction time curves for each monkey separately using Pearson correlations computed across *N* = 20 independent simulation runs differing in random seed, yielding stable estimates of mean model behavior and their associated standard errors. One-sample *t*-tests were used to assess whether mean Pearson correlations across seeds differed significantly from zero. To correct for multiple comparisons across the ten model configurations tested, the Benjamini-Hochberg False Discovery Rate procedure was applied at *q <* 0.05.

### 2.11 Explore-Exploit Trade-off

To quantify how quickly subjects abandon outdated value information following a target color switch, we measured perseveration to the previous block’s target color. For each trial, we calculated whether the chosen stimulus matched the optimal choice as determined by the preceding block’s target color, yielding a binary perseveration signal. Averaging this signal across all blocks produced a block-aligned perseveration curve as a function of trial position within the block (See Figure 6).

**Figure 6:**
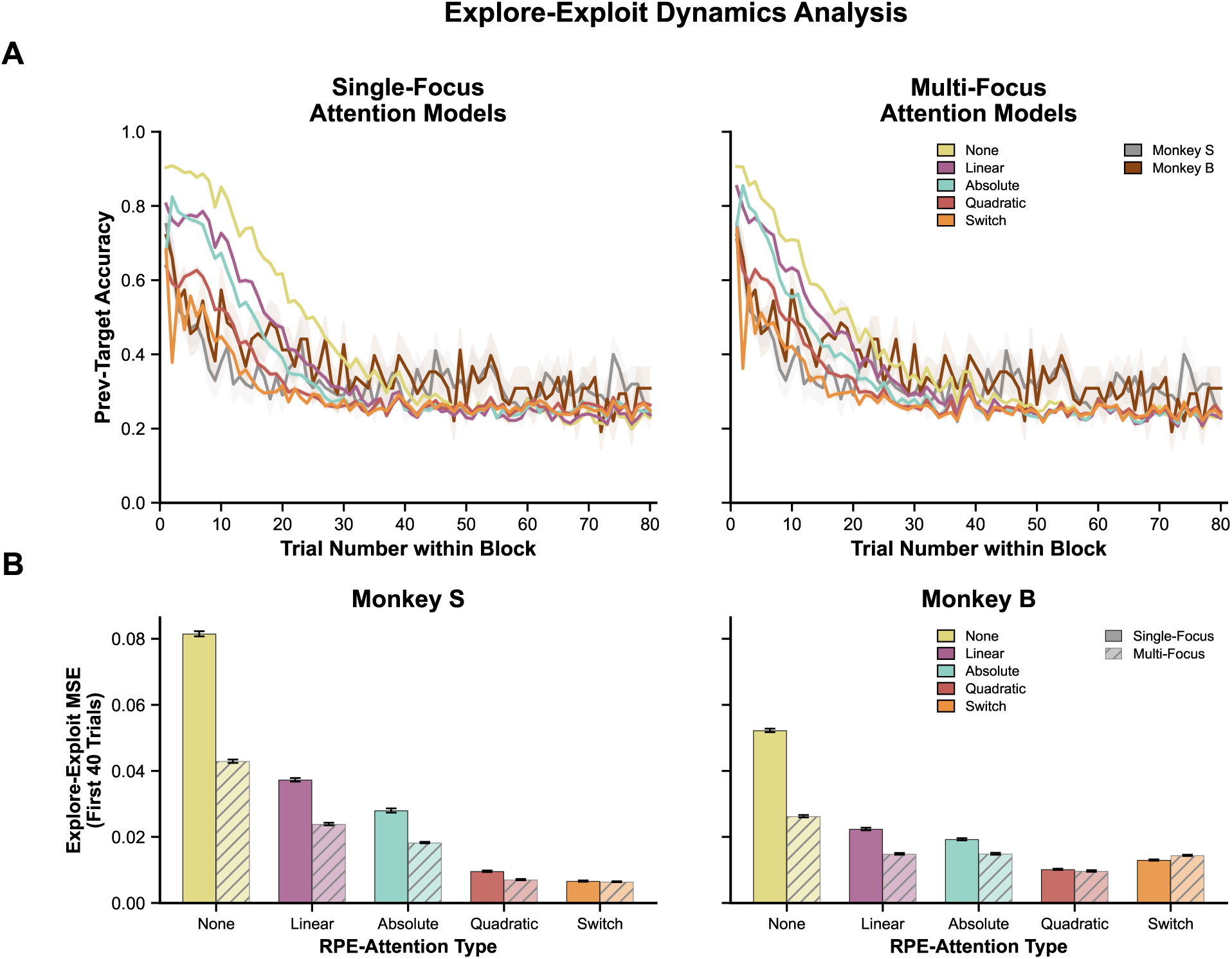
Explore-Exploit dynamics following target color switches. **(A)** Block-averaged accuracy with respect to the previous block’s target color as a function of trial number following a target color switch, for single-focus (left) and multi-focus (right) attention models. This metric indexes the explore-exploit trade-off: high accuracy on the previous target indicates continued exploitation of outdated value information, whereas rapid decay toward chance (33%) indicates a transition to exploration of the new reward landscape. Colored lines indicate model configurations defined by RPE-attention type (None, Linear, Absolute, Quadratic, Switch); solid dark lines indicate empirical data from Monkey B (brown) and Monkey S (gray). Model curves represent the mean across *N* = 20 independent simulation runs; monkey curves represent the mean across all valid blocks (excluding each monkey’s first block). Shaded regions denote ±1 SEM (across runs for models; across blocks for monkeys). **(B)** Mean squared error between each model’s exploration curve and the empirical monkey curve, computed over the first 40 trials following each target color switch, shown separately for Monkey B (left) and Monkey S (right). Bars indicate mean MSE across *N* = 20 runs; error bars indicate ±1 SEM. Solid bars, single-focus models; hatched bars, multi-focus models. Lower MSE indicates closer agreement with the empirical explore-exploit dynamics.

To compare model and monkey perseveration dynamics quantitatively, we computed the Mean Squared Error (MSE) between each model’s block-averaged perseveration curve and the corresponding empirical monkey curve over the first 40 trials following each target color switch. This window was chosen to capture the critical transient period during which perseveration to the previous target decays as the new target color is acquired. MSE was computed separately for each of the *N* = 20 independent simulation runs, and the mean and standard error of the mean were reported across runs for each model configuration and each monkey.

To rigorously quantify the speed at which subjects and computational models transitioned from exploration to exploitation of color values following a target color switch, we fitted an exponential decay function to the trial-by-trial perseveration curves. For both the empirical data and each independent model simulation run, we isolated the first 40 trials following a block transition and fitted the following single exponential decay function using non-linear least squares optimization:

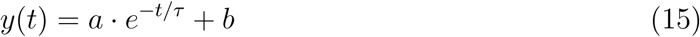

where *t* is the trial number following a block switch (1 ≤ *t* ≤ 40), *a* represents the initial amplitude of explore-exploit dynamic, *b* represents the asymptotic exploitation baseline, and *τ* is the decay time constant.

The parameter of interest was *τ*, which serves as an inverse index of learning speed; lower *τ* values indicate a more rapid abandonment of the outdated feature values and faster exploration of the new reward landscape. To ensure biologically and mathematically plausible fits, parameters were bounded during optimization (0 ≤ *a* ≤ 1.5; 0.5 ≤ *τ* ≤ 80; 0 ≤ *b* ≤ 1.0).

To statistically compare the exploration speed between model architectures, we computed *τ* for each independent simulation seed (*N* = 20). Fits that failed to explain a meaningful portion of the variance (*R*^2^ ≤ 0.3) were excluded to ensure that parameter comparisons were driven only by valid decay trajectories. We then performed pairwise Wilcoxon signed-rank tests on the distribution of *τ* values between models. To strictly control for multiple comparisons, all resulting *p*-values were adjusted using the Benjamini-Hochberg False Discovery Rate (FDR) procedure (See Figure 6).

### 2.12 Single Neuron Correlation

We performed single neuron correlations using neural data from Jahn et al. (2024). During monkey task performance, neural data was simultaneously recorded from three brain regions: lateral intraparietal area (LIP), frontal eye field (FEF), and prefrontal cortex (PFC).

#### Correlation Analysis

We computed Pearson correlations between individual neuron firing rates and previous trial RPE values across all recorded sessions. RPE values were obtained from the open-loop Q-learning behavioral model fit to each monkey’s choice behavior in Jahn et al. (2024). In the open-loop model, RPEs are calculated from the monkey’s choice on each trial and are therefore independent of the perceptual front-end and attentional modulation used for action selection in the present study.

Neuron firing rates were organized in 200-millisecond bins, aligned to trial stimulus onset. Neurons recorded on different recording days were treated as distinct neurons. We reported the firing rate for a bin at the bin midpoint (For example, activity recorded from 100 milliseconds before stimulus onset to 100 milliseconds after stimulus onset would be binned into the 0 millisecond time bin). Each neuron’s trial-by-trial activity was z-scored, and neurons with zero variance or missing values were excluded.

#### Shuffling Controls

To test for significance of the correlation, we randomly permute RPE values such that they are no longer chronologically sequenced while preserving the neural data temporal sequence.

#### Multiple Comparisons Correction

We used *p <* 0.05 to determine if a neuron had a significant correlation with RPE. We applied False Discovery Rate (FDR) correction using the Benjamini-Hochberg method (*q <* 0.05). FDR correction testing is applied to both real and shuffled data.

#### Effect Size

We quantified effect sizes using Cohen’s d and compared absolute correlation magnitudes between real and shuffled data using Mann-Whitney U tests.

All analyses were performed separately for each brain region (LIP, FEF, PFC), enabling comparison of RPE representation strength across different cortical areas involved in attention and learning. Correlation results were combined across the two monkeys (see Figure 7)

**Figure 7:**
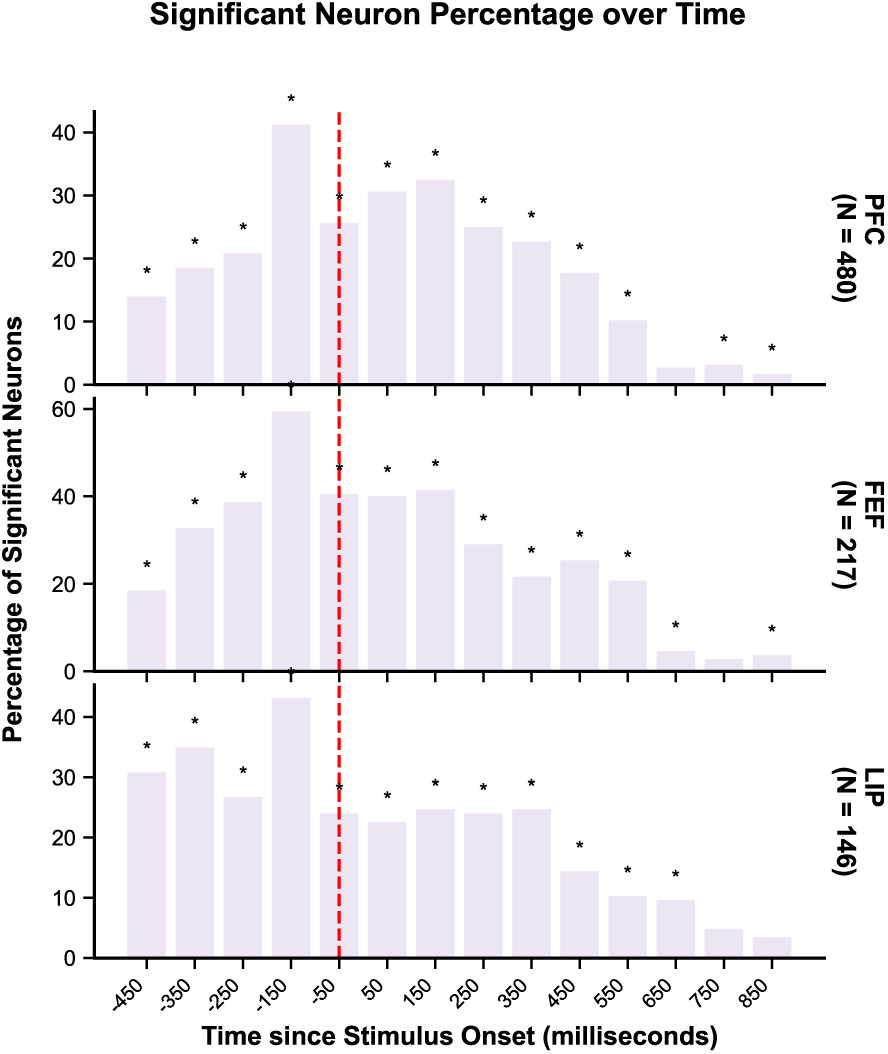
Percentage of neurons significantly correlated with previous-trial RPE across time. Percentage of neurons whose firing rate was significantly correlated with previous-trial RPE (Pearson correlation, *p <* 0.05, FDR-corrected within each recording session using Benjamini-Hochberg at *q <* 0.05), shown for PFC (top; *N* = 480 neurons), FEF (middle; *N* = 217), and LIP (bottom; *N* = 146). Firing rates were binned in 200 ms windows and aligned to current-trial stimulus onset (red dashed line at 0 ms). Each bar represents the percentage of neurons reaching significance in that time bin. Asterisks indicate time bins in which the proportion of significant neurons in the real data exceeded that in time-shuffled controls (Wilcoxon signed-rank test across sessions, FDR-corrected at *q <* 0.05). RPE values were derived from an open-loop baseline model without perceptual front-end or attentional modulation, using reward feedback based on each monkey’s empirical choices (Jahn et al., 2024).

### 2.13 Experimental Design and Statistical Analysis

Behavioral and neural data were collected by (Jahn et al., 2024) from two adult male rhesus macaques (*Macaca mulatta*); Monkey B, 13 kg; Monkey S, 9 kg). All experimental procedures were approved by the Princeton University Institutional Animal Care and Use Committee and conducted in accordance with NIH guidelines. Monkeys performed a color-value learning task across 17 recording sessions (8 sessions for Monkey B, 9 for Monkey S), yielding a total of 29,874 trials. Neural activity was recorded simultaneously from prefrontal cortex (PFC; 480 neurons), frontal eye fields (FEF; 217 neurons), and lateral intraparietal area (LIP; 146 neurons), totaling 843 neurons across both animals. (Jahn et al., 2024) recorded 492, 231, and 167 neurons from these regions, respectively; the reduced counts here reflect exclusion of neurons with missing values across the analyzed time windows.

For computational modeling, we simulated each of 10 model configurations (5 RPE-attention transfer functions × 2 attentional focus architectures) across *N* = 20 independent runs per configuration, each differing only in random seed, which impacts the stochastic action selection mechanism.

As described, pairwise comparisons between model configurations used Wilcoxon signed-rank tests on matched simulation runs, with all resulting *p*-values corrected for multiple comparisons using the Benjamini-Hochberg False Discovery Rate (FDR) procedure at *q <* 0.05. Ten pairwise comparisons were conducted per monkey for the learning curve analysis. Pearson correlations between block-averaged model decision entropy and empirical reaction time were computed separately for each simulation run; one-sample *t*-tests assessed whether the mean correlation across *N* = 20 runs differed significantly from zero, with FDR correction across the 10 model configurations tested per monkey.

To quantify explore-exploit dynamics, as mentioned, we fitted exponential decay functions (*y*(*t*) = *a* · *e*^−^*^t/τ^* + *b*) to trial-wise perseveration curves using nonlinear least squares, with parameter bounds 0 ≤ *a* ≤ 1.5, 0.5 ≤ *τ* ≤ 80, and 0 ≤ *b* ≤ 1.0. Fits with *R*^2^ ≤ 0.3 were excluded. Pairwise Wilcoxon signed-rank tests on the resulting *τ* distributions were corrected using BH-FDR at *q <* 0.05.

For the single-neuron correlation analysis, Pearson correlations were computed between individual neuron firing rates (in 200 ms bins aligned to stimulus onset) and previous-trial RPE values derived from the open-loop Q-learning model of (Jahn et al., 2024). Significance was assessed at *p <* 0.05, followed by FDR correction within each recording session at *q <* 0.05. Effect sizes were quantified using Cohen’s *d*, and absolute correlation magnitudes between real and shuffled data were compared using Mann-Whitney *U* tests. All shuffling controls preserved the temporal structure of neural data while randomly permuting RPE values.

Behavioral similarity analyses (entropy, maximal distance, minimal distance, mean distance) binned trial-wise accuracy into 20 equally spaced bins, with MSE computed between model and monkey accuracy curves across bins. All simulations and analyses were performed in Python using NumPy, SciPy, and Matplotlib.

### 2.14 Code Accessibility

The simulation and analysis code for this study is available upon request and will be openly available on GitHub upon publication. The behavioral and neural data from Jahn et al. (2024) are publicly available at https://doi.org/10.5281/zenodo.10529801.

## 3 Results

### 3.1 Rapid Acquisition and Sub-Optimal Plateaus in Monkey and Model Learning Trajectories

We first characterized the learning trajectories of the macaques in the color-value learning task (See Figure 1). Following a target color switch, monkeys displayed a characteristic bi-phasic learning profile: a rapid initial acquisition phase, achieving approximately 50% accuracy within 10–15 trials (chance level 33%), followed by a premature plateau at 75–80% accuracy. This asymptotic performance remained well below the theoretical optimum (100%), indicating persistent sub-optimal fine-grained color selection.

We then compared the ability of each computational architecture (Figure 2) to reproduce this specific bi-phasic temporal dynamic. Consistent with the fast timescale of monkey responses (∼220ms), our basic model architecture consists of bottom-up visually tuned responses gain-modulated by top-down attention, the result of which drives a priority map for saccade selection (Figure 3). Models varied along two dimensions: (1) the type of RPE-guided attention mechanism (None, Linear, Absolute, Quadratic, Switch; see Section 4.5), and (2) whether attention was allocated to a single color or distributed across multiple colors (single vs. multi; see Section 4.4).

All models demonstrate learning in this closed loop setting. This verifies that, as expected, none of the attention mechanisms explored here cause a negative feedback loop wherein misguided or rigid attention prevents exploration of value options. However, the amount and shape of learning differs across the different models.

To quantify model fit, we computed Mean Squared Error (MSE) between block-averaged model and monkey learning curves separately for each monkey (Figure 3B). Pairwise comparisons between single-focus models confirmed a hierarchical ordering of RPE-attention functions (Wilcoxon signed-rank tests, FDR-corrected at *q <* 0.05; Table 1). For both monkeys, each adjacent pair in the hierarchy None *>* Linear *>* Absolute *>* Quadratic/Switch differed significantly (all *p*_FDR_ *<* 0.001). The single-focus Quadratic and Switch models did not differ for Monkey B (*W* = 101, *p*_FDR_ = 0.898), but the Switch model yielded significantly lower MSE than the Quadratic model for Monkey S(*W* = 0, *p*_FDR_ = 1.907 × 10^−6^).

**Table 1:** Pairwise comparisons between single-focus model architectures on learning curve MSE. Wilcoxon signed-rank tests comparing mean squared error (MSE) between block-averaged model and monkey learning curves for all pairs of single-focus RPE-attention models. *p*-values were corrected for multiple comparisons using the Benjamini-Hochberg FDR procedure across 10 comparisons per monkey. *N* = 20 independent simulation runs per model configuration. \*\*\**p*_FDR_ *<* 0.001; ns, not significant (*p*_FDR_ ≥ 0.05).

| Comparison | Monkey B |  |  |  | Monkey S |  |  |  |
| --- | --- | --- | --- | --- | --- | --- | --- | --- |
| | $W$ | $p_{\text{FDR}}$ | Direction | | $W$ | $p_{\text{FDR}}$ | Direction | |
| None vs. Linear | 0 | $2.725 \times 10^{-6}$ | > | *** | 0 | $1.907 \times 10^{-6}$ | > | *** |
| None vs. Absolute | 0 | $2.725 \times 10^{-6}$ | > | *** | 0 | $1.907 \times 10^{-6}$ | > | *** |
| None vs. Quadratic | 0 | $2.725 \times 10^{-6}$ | > | *** | 0 | $1.907 \times 10^{-6}$ | > | *** |
| None vs. Switch | 0 | $2.725 \times 10^{-6}$ | > | *** | 0 | $1.907 \times 10^{-6}$ | > | *** |
| Linear vs. Absolute | 0 | $2.725 \times 10^{-6}$ | > | *** | 0 | $1.907 \times 10^{-6}$ | > | *** |
| Linear vs. Quadratic | 0 | $2.725 \times 10^{-6}$ | > | *** | 0 | $1.907 \times 10^{-6}$ | > | *** |
| Linear vs. Switch | 0 | $2.725 \times 10^{-6}$ | > | *** | 0 | $1.907 \times 10^{-6}$ | > | *** |
| Absolute vs. Quadratic | 14 | $2.623 \times 10^{-4}$ | > | *** | 0 | $1.907 \times 10^{-6}$ | > | *** |
| Absolute vs. Switch | 20 | $7.863 \times 10^{-4}$ | > | *** | 0 | $1.907 \times 10^{-6}$ | > | *** |
| Quadratic vs. Switch | 101 | $8.983 \times 10^{-1}$ | < | ns | 0 | $1.907 \times 10^{-6}$ | > | *** |

Direction indicates which model had higher median MSE (i.e., worse fit): “>” means the first-listed model had higher MSE than the second. For the Quadratic vs. Switch comparison in Monkey B, “<” indicates Quadratic had slightly lower median MSE, but the difference was not significant.

A difference in model ranking for the two monkeys is perhaps not surprising, given that the two monkeys differed in both task parameters and learning dynamics. Monkey B faced a lower accuracy threshold for target switches (80% vs. 85% for Monkey S) and received an escalating reward schedule, whereas Monkey S received fixed rewards (Jahn et al., 2024). Despite the lower threshold, Monkey B required more trials to trigger switches, consistent with slower value learning. Monkey S was the more competent learner of the pair, showing clearer and more consistent behavioral patterns across analyses. Regarding attentional focus, single-focus architectures yielded significantly lower MSE than their multi-focus counterparts for all RPE-modulated models (Linear, Absolute, Quadratic, Switch; all *p*_FDR_ *<* 0.004; Table 2). The sole exception was the None condition, where multi-focus models produced lower MSE than single-focus models for both monkeys (Monkey B: *p*_FDR_ = 3.338×10^−5^; Monkey S: *p*_FDR_ = 4.768 × 10^−6^), indicating that distributing attention across features is advantageous only in the absence of RPE-guided modulation.

**Table 2:** Single-focus vs. multi-focus attention architecture comparisons on learning curve MSE. Wilcoxon signed-rank tests comparing MSE between single-focus and multi-focus variants of each RPE-attention model. *p*-values were corrected using Benjamini–Hochberg FDR across 5 comparisons per monkey. *N* = 20 independent simulation runs per configuration. \*\**p*_FDR_ *<* 0.01; \*\*\**p*_FDR_ *<* 0.001.

| RPE Type | Monkey B |  |  |  | Monkey S |  |  |  |
| --- | --- | --- | --- | --- | --- | --- | --- | --- |
| | $W$ | $p_{\text{FDR}}$ | Winner | | $W$ | $p_{\text{FDR}}$ | Winner | |
| None | 6 | $3.338 \times 10^{-5}$ | Multi | *** | 0 | $4.768 \times 10^{-6}$ | Multi | *** |
| Linear | 29 | $3.153 \times 10^{-3}$ | Single | ** | 27 | $2.325 \times 10^{-3}$ | Single | ** |
| Absolute | 0 | $9.537 \times 10^{-6}$ | Single | *** | 2 | $9.537 \times 10^{-6}$ | Single | *** |
| Quadratic | 1 | $9.537 \times 10^{-6}$ | Single | *** | 11 | $1.311 \times 10^{-4}$ | Single | *** |
| Switch | 2 | $9.537 \times 10^{-6}$ | Single | *** | 0 | $4.768 \times 10^{-6}$ | Single | *** |

“Winner” indicates the architecture with lower median MSE (better fit). For the None condition, the multi-focus architecture provided a better fit in both monkeys. For all RPE-modulated conditions, the single-focus architecture was superior.

In total, we observe that the single-focused Switch model provides the best fit when considering both monkeys and is able to capture a fast rise and sub-optimal plateau in performance. Notably, these features of the learning curve cannot be explained by simply adjusting the learning rate parameter; merely varying learning rate fails to simultaneously capture both the rapid initial acquisition and the low asymptotic accuracy observed in the monkeys (see Extended Data Figure 9).

### 3.2 Single-Focus Attention Architectures Consistently Outperform Multi-Focus Models Across Behavioral Similarity Metrics

The block-averaged learning curve captures whether models achieve similar overall accuracy to the monkeys over the course of learning, but accuracy is a single, coarse-grained measure. Many different patterns of correct and incorrect choices can produce the same average accuracy. We therefore developed a complementary set of novel behavioral similarity analyses, parameterizing individual trials according to four stimulus-configuration metrics that capture distinct aspects of trial difficulty and examining how accuracy varied as a function of each metric for both models and monkeys (see Section 4.9).

#### Entropy

Monkey behavioral data revealed a robust inverse relationship between stimulus value entropy and accuracy: performance peaked on low-entropy trials (accuracy ≈ 0.90) and degraded monotonically as the value distribution became more uniform, dropping to ≈ 0.45 at maximal entropy, confirming that monkeys struggled to discriminate targets when stimulus values were approximately equal. All models showed the same overall shape of this relationship but had different quantitative fits. Single-focus architectures produced lower MSE than their multi-focus counterparts for most RPE conditions (Table 5). Among single-focus models, the Absolute model achieved the lowest MSE for Monkey B, significantly outperforming its multi-focus equivalent (Wilcoxon signed-rank, *W* = 0, *p*_FDR_ *<* 1.0 × 10^−4^). For Monkey S, the Switch model ranked first. However, direct pairwise comparisons between the top two single-focus models (Single Absolute vs. Single Switch) revealed no significant difference in fit for either Monkey B (*W* = 54, *p*_FDR_ = 0.065) or Monkey S (*W* = 61, *p*_FDR_ = 0.105; Table 3).

**Table 3:** Pairwise comparisons between single-focus models on the entropy behavioral similarity metric. Wilcoxon signed-rank tests comparing MSE between binned model and monkey accuracy curves as a function of trial-wise stimulus entropy. *p*-values were corrected using Benjamini–Hochberg FDR across 10 comparisons per monkey. *N* = 20 simulation runs per configuration. \**p*_FDR_ *<* 0.05; \*\**p*_FDR_ *<* 0.01; \*\*\**p*_FDR_ *<* 0.001; ns, not significant.

| Comparison | Monkey B |  |  | Monkey S |  |  |
| --- | --- | --- | --- | --- | --- | --- |
| | $W$ | $p_{\text{FDR}}$ | | $W$ | $p_{\text{FDR}}$ | |
| None vs. Linear | 0 | $<10^{-4}$ | *** | 0 | $<10^{-4}$ | *** |
| None vs. Absolute | 0 | $<10^{-4}$ | *** | 0 | $<10^{-4}$ | *** |
| None vs. Quadratic | 0 | $<10^{-4}$ | *** | 0 | $<10^{-4}$ | *** |
| None vs. Switch | 0 | $<10^{-4}$ | *** | 0 | $<10^{-4}$ | *** |
| Linear vs. Absolute | 0 | $<10^{-4}$ | *** | 0 | $<10^{-4}$ | *** |
| Linear vs. Quadratic | 0 | $<10^{-4}$ | *** | 43 | 0.021 | * |
| Linear vs. Switch | 2 | $<10^{-4}$ | *** | 0 | $<10^{-4}$ | *** |
| Absolute vs. Quadratic | 35 | 0.009 | ** | 14 | $3.0 \times 10^{-4}$ | *** |
| Absolute vs. Switch | 54 | 0.065 | ns | 61 | 0.105 | ns |
| Quadratic vs. Switch | 86 | 0.498 | ns | 5 | $<10^{-4}$ | *** |

#### Maximal Distance

Monkey accuracy increased as the angular distance between the target and the furthest available stimulus grew from 0 to ≈ 2.2 radians, with accuracy rising from 0.55 to 0.62, before saturating at ≈ 0.64 for distances approaching *π* radians. This is consistent with the notion that having at least one stimulus of low value eases the decision. For the models, single-focus attention achieved closer fits to both monkey curves than multi-focus counterparts across most RPE conditions, with the exception of the Quadratic condition for both monkeys, where the multi-focus Quadratic model outperformed its single-focus equivalent. Among single-focus models, the Absolute model ranked first and Switch ranked second for both Monkey B and Monkey S. Once again, the difference between the Single Absolute and Single Switch models was not statistically significant for either monkey (Monkey B: *W* = 85, *p*_FDR_ = 0.475; Monkey S: *W* = 85, *p*_FDR_ = 0.475).

#### Minimal Distance

Monkeys show the opposite performance pattern for minimal distance as for maximal, demonstrating how having at least one high value stimulus eases the decision. Among single-focus models, the Quadratic model achieved the lowest MSE for Monkey B, significantly outperforming Multi Quadratic (*W* = 4, *p*_FDR_ *<* 1.0 × 10^−4^; Table 6), while Switch ranked second (*W* = 97, *p*_FDR_ = 0.870 vs. Quadratic; Table 4). For Monkey S, the Single Switch model ranked first, yielding significantly lower MSE than Multi Switch (*W* = 19, *p*_FDR_ = 7.0 × 10^−4^), while the Single Absolute ranked second (*W* = 75, *p*_FDR_ = 0.277 vs. Switch).

**Table 4:** Pairwise comparisons between single-focus models on the minimal distance behavioral similarity metric. Wilcoxon signed-rank tests comparing MSE between binned model and monkey accuracy curves as a function of minimal stimulus–target angular distance. *p*-values were corrected using Benjamini–Hochberg FDR across 10 comparisons per monkey. *N* = 20 simulation runs per configuration. \*\**p*_FDR_ *<* 0.01; \*\*\**p*_FDR_ *<* 0.001; ns, not significant.

| Comparison | Monkey B |  |  | Monkey S |  |  |
| --- | --- | --- | --- | --- | --- | --- |
| | $W$ | $p_{\text{FDR}}$ | | $W$ | $p_{\text{FDR}}$ | |
| None vs. Linear | 0 | $<10^{-4}$ | *** | 0 | $<10^{-4}$ | *** |
| None vs. Absolute | 0 | $<10^{-4}$ | *** | 0 | $<10^{-4}$ | *** |
| None vs. Quadratic | 0 | $<10^{-4}$ | *** | 0 | $<10^{-4}$ | *** |
| None vs. Switch | 0 | $<10^{-4}$ | *** | 0 | $<10^{-4}$ | *** |
| Linear vs. Absolute | 0 | $<10^{-4}$ | *** | 1 | $<10^{-4}$ | *** |
| Linear vs. Quadratic | 0 | $<10^{-4}$ | *** | 0 | $<10^{-4}$ | *** |
| Linear vs. Switch | 0 | $<10^{-4}$ | *** | 0 | $<10^{-4}$ | *** |
| Absolute vs. Quadratic | 97 | 0.870 | ns | 62 | 0.127 | ns |
| Absolute vs. Switch | 100 | 0.870 | ns | 75 | 0.277 | ns |
| Quadratic vs. Switch | 97 | 0.870 | ns | 25 | 0.002 | ** |

**Table 5:**
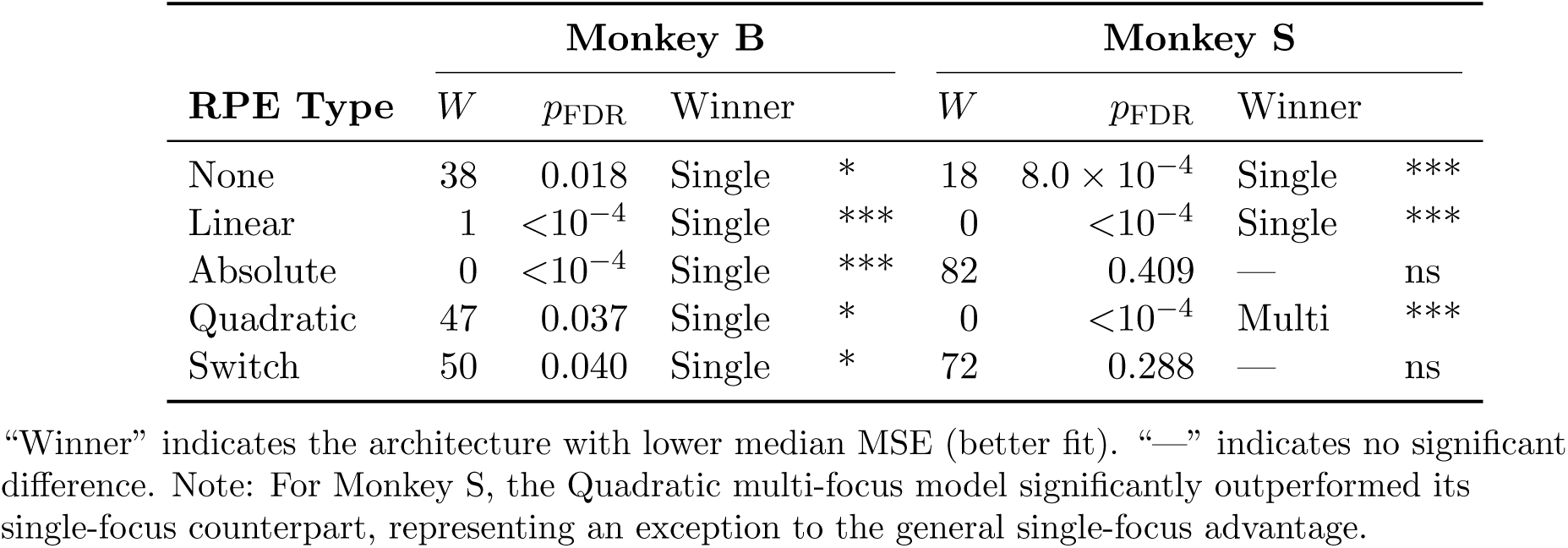
Single-focus vs. multi-focus comparisons on the entropy behavioral similarity metric. Wilcoxon signed-rank tests comparing MSE between single-focus and multi-focus variants. *p*-values corrected using Benjamini-Hochberg FDR across 5 comparisons per monkey. *N* = 20 simulation runs. \**p*_FDR_ *<* 0.05; \*\*\**p*_FDR_ *<* 0.001; ns, not significant.

**Table 6:** Single-focus vs. multi-focus comparisons on the minimal distance behavioral similarity metric. Wilcoxon signed-rank tests comparing MSE between single-focus and multi-focus variants. *p*-values corrected using Benjamini–Hochberg FDR across 5 comparisons per monkey. Note: For Monkey S Quadratic, the multi-focus architecture yielded lower MSE than single-focus (*p*_FDR_ = 0.004). *N* = 20 simulation runs. \*\**p*_FDR_ *<* 0.01; \*\*\**p*_FDR_ *<* 0.001.

| RPE Type | Monkey B |  |  |  | Monkey S |  |  |  |
| --- | --- | --- | --- | --- | --- | --- | --- | --- |
| | $W$ | $p_{\text{FDR}}$ | Winner | | $W$ | $p_{\text{FDR}}$ | Winner | |
| None | 4 | $<10^{-4}$ | Single | *** | 6 | $1.0 \times 10^{-4}$ | Single | *** |
| Linear | 0 | $<10^{-4}$ | Single | *** | 0 | $<10^{-4}$ | Single | *** |
| Absolute | 0 | $<10^{-4}$ | Single | *** | 7 | $1.0 \times 10^{-4}$ | Single | *** |
| Quadratic | 4 | $<10^{-4}$ | Single | *** | 31 | $4.2 \times 10^{-3}$ | Multi | ** |
| Switch | 2 | $<10^{-4}$ | Single | *** | 19 | $7.0 \times 10^{-4}$ | Single | *** |

#### Mean Distance

Monkey accuracy rose from ≈ 0.50 at the smallest mean distances, plateaued near ≈ 0.70 at a mean distance of ≈ 1.8 radians, and subsequently declined at larger distances. For Monkey B, the Single Absolute model produced the lowest MSE, significantly outperforming Multi Absolute (*W* = 19, *p*_FDR_ = 1.0 × 10^−3^). For Monkey S, Single Switch ranked first, significantly outperforming Multi Switch (*W* = 8, *p*_FDR_ = 1.0 × 10^−4^). Consistent with the other metrics, the top two single-focus models did not differ significantly from one another for Monkey B (Absolute vs. Quadratic, *W* = 88, *p*_FDR_ = 0.607). For Monkey S, the Single Switch model yielded significantly lower MSE than the Single Absolute model (Switch vs. Absolute, *W* = 45, *p*_FDR_ = 0.027).

As we can see, across all four trial difficulty metrics, single-focus attention architectures consistently provide a better match to monkey data, in part due to the over performance of the multi-focus models on easy trials. This likely results from the ability of multi-focus to distribute attentional enhancement and suppression more freely across stimuli of different values.

### 3.3 Model Confidence Approximates Empirical Reaction Time Dynamics

We applied a novel analysis of reaction time to infer monkey confidence over learning. For Monkey S, the stronger learner of the pair, mean RT rose from approximately 215 ms during the first 10 trials to 224 ms during the latter half of the block (trials 41 − 80), showing a clear and consistent upward trend. Monkey B showed a similar pattern, though weaker, with mean RT rising from approximately 204 ms during the first 10 trials to 212 ms over the same intervals. In both animals, this increase was concentrated in the first 40 trials (linear slope: Monkey B: 0.23 ms/trial, *p* = 0.024; Monkey S: 0.26 ms/trial, *p* = 0.002) before plateauing in the second half. This increase in reaction time during early learning is perhaps surprising, as one might expect faster responses to accompany improved accuracy.

We therefore asked which model architectures produced decision entropy dynamics that tracked this empirical RT pattern. We used the Shannon entropy of each model’s choice probability distribution as a proxy for decision confidence (see Section 4.10), reasoning that higher entropy (greater choice uncertainty) should correspond to slower responses. We then asked which model architectures produced entropy trajectories that tracked the empirical RT pattern.

Through inspection of Figure 5, we can see qualitative differences in the models with different RPE modulation, with only Absolute and Switch models showing the necessary increase and Linear and Quadratic showing a decrease. This is supported by quantitative analysis. Across both monkeys, Pearson correlations between block-averaged model entropy and empirical reaction times showed only the Absolute and Switch RPE-guided attention models yielded consistently positive correlations (single-focus Absolute: Monkey B: *r* = 0.31, Monkey S: *r* = 0.39; single-focus Switch: Monkey B: *r* = 0.17, Monkey S: *r* = 0.42; all *p*_FDR_ *<* 0.001; Table 7). While most models reached statistical significance after FDR correction due to the large number of simulation runs, the None, Linear, and Quadratic models produced near-zero or negative correlations for at least one monkey, failing to capture the expected positive relationship between model uncertainty and empirical response latency. Direct pairwise comparisons confirmed this advantage: both the Single Absolute and Single Switch models yielded significantly higher correlations than the Single Quadratic model across both subjects (Wilcoxon signed-rank, all *p*_FDR_ *<* 0.001; Table 8). See Figure 5.

**Table 7:** Mean Pearson correlations between model decision entropy and empirical reaction time. One-sample *t*-tests assessed whether the mean Pearson *r* across *N* = 20 simulation runs differed significantly from zero. *p*-values were corrected using Benjamini–Hochberg FDR across 10 model configurations per monkey. Values shown are mean *r*± SEM. \*\**p*_FDR_ *<* 0.01; \*\*\**p*_FDR_ *<* 0.001; ns, not significant.

| RPE Type | Focus | Monkey B |  |  | Monkey S |  |  |
| --- | --- | --- | --- | --- | --- | --- | --- |
| | | $\bar{r} \pm \text{SEM}$ | $p_{\text{FDR}}$ | | $\bar{r} \pm \text{SEM}$ | $p_{\text{FDR}}$ | |
| None | Single | $+0.041 \pm 0.010$ | $5.93 \times 10^{-4}$ | *** | $+0.190 \pm 0.006$ | $5.79 \times 10^{-18}$ | *** |
| None | Multi | $-0.043 \pm 0.012$ | $2.06 \times 10^{-3}$ | ** | $-0.085 \pm 0.010$ | $1.59 \times 10^{-7}$ | *** |
| Linear | Single | $+0.039 \pm 0.006$ | $4.73 \times 10^{-6}$ | *** | $-0.233 \pm 0.006$ | $8.07 \times 10^{-19}$ | *** |
| Linear | Multi | $-0.097 \pm 0.011$ | $4.83 \times 10^{-8}$ | *** | $-0.414 \pm 0.005$ | $2.74 \times 10^{-25}$ | *** |
| Absolute | Single | $+0.311 \pm 0.003$ | $5.97 \times 10^{-27}$ | *** | $+0.392 \pm 0.003$ | $9.72 \times 10^{-30}$ | *** |
| Absolute | Multi | $+0.231 \pm 0.007$ | $2.49 \times 10^{-17}$ | *** | $+0.198 \pm 0.008$ | $6.51 \times 10^{-16}$ | *** |
| Quadratic | Single | $+0.126 \pm 0.007$ | $6.36 \times 10^{-13}$ | *** | $-0.081 \pm 0.009$ | $5.71 \times 10^{-8}$ | *** |
| Quadratic | Multi | $-0.151 \pm 0.006$ | $3.21 \times 10^{-15}$ | *** | $-0.469 \pm 0.005$ | $2.44 \times 10^{-26}$ | *** |
| Switch | Single | $+0.170 \pm 0.008$ | $4.84 \times 10^{-14}$ | *** | $+0.421 \pm 0.008$ | $3.79 \times 10^{-21}$ | *** |
| Switch | Multi | $+0.003 \pm 0.010$ | $7.46 \times 10^{-1}$ | ns | $+0.173 \pm 0.009$ | $1.19 \times 10^{-13}$ | *** |

**Table 8:**
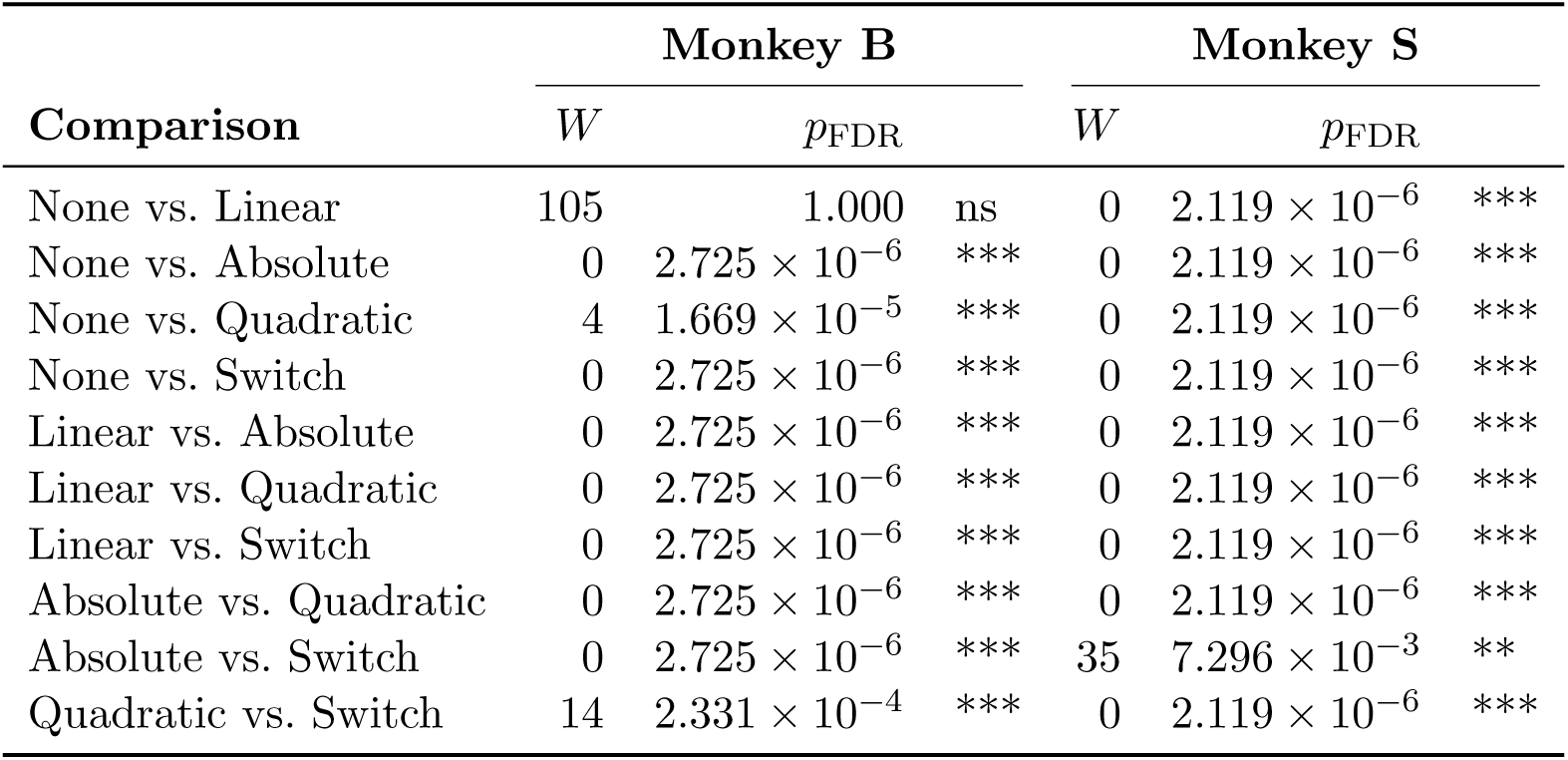
Pairwise comparisons between single-focus models on entropy–reaction time correlation. Wilcoxon signed-rank tests comparing Pearson *r* values between model configurations. *p*-values corrected using Benjamini-Hochberg FDR across 10 comparisons per monkey. *N* = 20 simulation runs. \*\*\**p*_FDR_ *<* 0.001; ns, not significant.

The qualitative differences in entropy trajectories across model types can be understood by examining the shape of each RPE-attention transfer function in the critical range of RPE values traversed during early learning. In the initial trials following a target switch, the model receives large, and predominantly negative, RPE as it selects stimuli based on the old target information. As learning proceeds, RPE values migrate toward zero. For the Absolute and Switch transfer functions, attention strength decreases as RPE moves from negative values toward zero (see Figure 5), producing rising entropy over trials as the attentional gain field weakens. In contrast, for the Linear and Quadratic transfer functions, attention strength increases in this same negative-to-zero RPE range, producing a decline in entropy that does not match the empirical RT increase. The None model, lacking any RPE modulation, produces a flat entropy trajectory that similarly fails to capture the observed RT dynamics. This mechanistic explanation accounts for why only a subset of the RPE-attention transfer functions yield entropy trajectories that positively correlate with empirical reaction time.

Within each RPE model type, single-focus attention architectures consistently produced significantly higher Pearson correlations than their multi-focus counterparts across both monkeys and all RPE conditions. Specifically, the Single Absolute model significantly out-performed the Multi-Absolute model (Monkey B: *W* = 0, *p*_FDR_ = 2.384 × 10^−6^; Monkey S: *W* = 0, *p*_FDR_ = 1.907 × 10^−6^), and the Single Switch model significantly outperformed the Multi Switch model (Monkey B: *W* = 0, *p*_FDR_ = 2.384 × 10^−6^; Monkey S: *W* = 0, *p*_FDR_ = 1.907 × 10^−6^). This pattern held without exception, indicating a reliable advantage of winner-take-all attentional selection over distributed attention for approximating behavioral response vigor.

The two monkeys showed partially overlapping but distinct rank orderings of model fit. For Monkey S, the single-focus Switch model produced the strongest positive correlation, slightly but significantly outperforming the single-focus Absolute model (*W* = 35, *p*_FDR_ = 7.296×10^−3^). For Monkey B, the single-focus Absolute model yielded the highest correlation, providing a significantly better fit to the reaction time data than the single-focus Switch model (*W* = 0, *p*_FDR_ = 2.725 × 10^−6^).

Taken together, single-focus Absolute and Switch RPE-guided attention models produced the strongest and most consistent positive correlations with empirical reaction time across both monkeys.

### 3.4 Explore-Exploit Trade-Off

A benefit of RPE-guided attention is that it can allow for corrective responses after errors. To assess how well each model captured the temporal dynamics of exploration following a target color switch, we examined block-averaged explore-exploit dynamic curves, that is, trial-wise accuracy with respect to the previous block’s target color, for both monkeys and all model configurations. (Figure 6)

Following a target color switch, both monkeys rapidly abandoned the previous block’s target color, with previous-target accuracy decaying from above 60% to near chance within approximately 15 − 20 trials, consistent with the rapid forgetting of previous templates reported in Jahn et al. (2024). Model analysis revealed that the None, Linear, and Absolute single-focus models failed to replicate the rapid decay in previous-target accuracy observed in both monkeys over the first 40 trials. These models exhibited persistently elevated perseveration, decaying more slowly than the empirical monkey curves.

Quantitative MSE analysis over the first 40 trials confirmed this pattern (Figure 6). Both the Single Switch and Single Quadratic models yielded significantly lower MSE than the Single Absolute model during these initial 40 trials (Wilcoxon signed-rank, all *p*_FDR_ ≤ 2.384 × 10^−6^ for both monkeys). Among the higher-MSE models, multi-focus architectures consistently produced lower MSE than their single-focus counterparts. However, for the lowest-MSE configurations (Quadratic for Monkey B and Switch for Monkey S) single- and multi-focus models yielded comparable MSE values. Across both monkeys, the Quadratic and Switch models achieved the lowest overall MSE. Direct comparison between these two top-performing models (i.e., Single Switch and Single Quadratic) revealed idiosyncratic advantages on this specific metric: the Single Quadratic model provided a significantly lower MSE for Monkey B (*W* = 14, *p*_FDR_ = 2.331 × 10^−4^), while the Single Switch model was significantly superior for Monkey S (*W* = 16, *p*_FDR_ = 3.223 × 10^−4^). For Monkey S, the Single Switch model produced the lowest MSE among all configurations, though it did not differ significantly from the Multi Switch model (Wilcoxon signed-rank, *W* = 95, *p*_FDR_ = 0.729).

A core prediction of the Switch mechanism is that inverting attentional gain following negative RPE should accelerate exploration of alternative feature values after a target switch. To test this prediction, we fitted an exponential decay function to the explore-exploit dynamic trajectories to extract the decay time constant (*τ*; see Section 4.11). Both monkeys exhibited highly rapid behavioral transitions away from the previous target (Monkey B: *τ* = 11.30; Monkey S: *τ* = 4.05). Consistent with this prediction, among the computational models, the Single Switch architecture produced the fastest exploration dynamics. Across simulation runs, the Single Switch model yielded a significantly faster (lower) decay time constant (mean *τ* ≈ 12.85) compared to the Single Quadratic model (mean *τ* ≈ 18.93; *W* = 0, *p*_FDR_ = 1.907 × 10^−6^) and the Single Absolute model (mean *τ* ≈ 23.41; *W* = 0, *p*_FDR_ = 1.907 × 10^−6^). The Single Switch model also decayed significantly faster than both the baseline without any RPE modulation (Single None, mean *τ* ≈ 33.78; *W* = 0, *p*_FDR_ = 1.907×10^−6^) and the Linear model (mean *τ* ≈ 29.98; *W* = 0, *p*_FDR_ = 1.907 × 10^−6^).

### 3.5 Neural Evidence Supporting RPE-Guided Attention

We have thus far found convincing behavioral evidence that previous-trial RPE is used to modulate attention. To investigate whether neural activity in attention-related brain regions carries RPE information consistent with such modulation, we analyzed simultaneously-recorded neural data while the monkey performed the color-value learning task (Jahn et al., 2024).

We computed Pearson correlations between individual neuron firing rates and previous-trial RPE values across all recorded sessions. RPE values were derived from the Q-learning behavioral model fit to each monkey’s choice behavior (Jahn et al., 2024)(see Section 4.12). Statistical significance was assessed at *p <* 0.05, followed by false discovery rate (FDR) correction to account for multiple comparisons across the 843 included neurons.

Neural activity was analyzed across 14 consecutive 200-millisecond time windows spanning −550 milliseconds to 950 milliseconds centered on current trial stimulus onset. After FDR correction, for PFC, 37.5% of neurons were significantly correlated to previous-trial RPE (180 out of 480). For FEF, 41.9% of neurons were significantly correlated to previous-trial RPE (91 out of 217). For LIP, 26.7% of neurons were significantly correlated to previous-trial RPE (39 out of 146) (see Figure 7).

RPE-correlated neural activity peaked in the time bin centered at −150 milliseconds relative to stimulus onset, a pattern consistent across all three brain regions (see Figure 7). At the peak time bin, individual correlation magnitudes were modest: median |*r*| = 0.086 for PFC (interquartile range: 0.074 - 0.105), 0.076 for FEF (0.067-0.092), and 0.081 for LIP (0.066-0.105). Such a time course of resurgence in RPE encoding directly before the onset of the search array of the next trial is consistent with the use of RPE in attentional modulation; attention has been shown to modulate activity in anticipation of sensory stimuli (Giesbrecht et al., 2006).

Correlated neurons showed a positive bias in correlation sign in FEF (71% positively correlated neurons) and LIP (83%). However PFC was more evenly split (54% positively correlated). To implement a switch-like mechanism, having equal populations of positively and negatively correlated neurons may be beneficial: neurons positively correlated with RPE can drive attention towards stimuli currently associated with high value while those negatively correlated with RPE can drive it toward stimuli associated with low value.

Furthermore, correlating neural activity with unsigned rather than signed RPE yields far fewer significant neurons (PFC: 7.1%, LIP: 9.1%, FEF: 8.7%). This suggests that absolute value of RPE may not be driving attentional modulation. Overall, our neural analysis lends support for the switch mechanism.

## 4 Discussion

We investigated the computational algorithms linking reinforcement learning to feature-based attention by analyzing monkey behavior and neural activity and building candidate mechanistic models. Through this, we conclude that a single-focus, RPE-modulated attention mechanism is most consistent with animal behavior. Specifically, our results demonstrate that the temporal dynamics of primate learning, particularly the rapid initial acquisition followed by a sub-optimal plateau, are best explained by a single-focus “Switch” attentional architecture. In this framework, attention operates as a winner-take-all filter that focuses resources on the highest-valued feature but transiently inverts to prioritize exploration following negative prediction errors.

Evidence in favor of the single switch mechanism is strongest for Monkey S, the better learner of the pair and comes in the form of a better match to learning dynamics, error patterns, confidence changes, and explore-exploit behaviors. Even where the single switch mechanism does not provide the best quantitative fit (especially for Monkey B) it usually is competitive with a different single-focus RPE-modulated model (either Absolute or Quadratic). Furthermore, our neural analysis suggests that PFC could support a switch-like mechanism.

Standard reinforcement learning models often assume that agents have uniform access to the state space. However, our findings regarding single vs. multi-focus attention indicate that biological learning is constrained by an attentional bottleneck. Interestingly, in most cases, the multi-focus attention models modulated choice probabilities too effectively, resulting in a much higher accuracy than that of the monkeys, especially in later trials in a block. The single-focus attention model captures a constraint that may be responsible for sub-optimal behavior. The task design required the use of covert attention: monkeys had to maintain central fixation during stimulus presentation (Jahn et al., 2024) before saccading to choose a color stimulus. Our finding aligns with established constraints on covert attention capacity as studies have shown that attention to multiple features simultaneously comes at a significant cost to processing efficiency (Desimone et al. (1995); Grubert and Eimer (2015); Liu and Jigo (2017); Grubert et al. (2016); though note Williams et al. (2023)).

While purely positive feedback loops (attending to what is valuable) facilitate exploitation, they carry the risk of perseveration. To protect against this, we hypothesized ways by which attention could be modulated by RPE feedback. Through this, we identified a specific transfer function, the “Switch” mechanism, that can facilitate rapid error-based exploration by using negative RPE to rapidly explore new value distributions. When an expected reward is not given, the negative RPE inverts the gain landscape, suppressing the previously attended feature and enhancing the salience of competing features. This mirrors the adaptive exploration hypothesis, where exploration is not random noise but a directed process triggered by unexpected negative outcomes (Sallet et al., 2013; Cohen et al., 2007).

Another relevant implication is that, as RPE weakens over learning, RPE-modulated attention strength weakens, decreasing the magnitude of attention. Because attention drives action selection, this weakening could account for both the suboptimal plateau in learning curves and the observed increase in reaction times. This aligns with Schultz et al. (1997)’s empirical finding that suggests RPE signals weaken as learning progresses and we connect this decrease in RPE signaling to a decrease in attention strength, which impacts performance. Furthermore, attention-related neural activity has also been shown to decrease over the course of perceptual learning, which could be the result of decreased RPEs (Mukai et al., 2007; Yotsumoto et al., 2008; Sigman et al., 2005). Additionally, a previous study found that the absence of reward on previous trials predicted switches of attention across stimulus dimensions (Leong et al., 2017). Our study focuses on switches of attention within a single feature dimension (color hue). Despite this difference in scope, both findings converge on the principle that negative prediction errors trigger attentional reorientation, supporting the generality of error-driven exploration mechanisms.

In total, the switch mechanism provides a normative explanation for the observed behavioral limits of the monkeys: the visual system sacrifices asymptotic accuracy to maximize the speed of value learning in volatile environments. Future work could explore how different environmental statistics (e.g. rate of target color change) impact the relationship between RPE and attention.

Our results build on prior work linking prediction errors to attentional allocation. Oemisch et al. (2019) identified feature-specific prediction error signals in macaque fronto-striatal circuits, and Torrents-Rodas et al. (2021) showed that prediction errors broaden attention to previously irrelevant cues. While these studies established that RPE can influence attention, they did not specify the mathematical form of this influence. Our modeling work addresses this gap by testing five candidate transfer functions and, while the Switch mechanism works best overall, we find that unsigned surprise (the Absolute mechanism) can also adequately capture primate learning dynamics. We can therefore narrow down the likely form that a function connecting RPE to attention strength takes to one where attention weakens as RPEs decrease in magnitude. Knowing this relationship can help interpret data about attentional modulation of neural activity in the visual system. Cross-trial variations that may have previously been considered noise may be attributable to fluctuations in previous trial RPE.

Our neural analyses further support the hypothesis that RPE signals modulate attentional processes. Single-neuron correlations revealed substantial proportions of neurons in PFC, FEF, and LIP regions whose firing rates tracked previous-trial RPE, with correlations peaking prior to the next stimulus onset. As reported in 5.5, however, individual RPE-firing rate correlations were small (median |*r*| ≈ 0.08 across regions). The neurons recorded by Jahn et al. (2024) were not targeted for RPE sensitivity and RPE is likely one of many variables encoded in these mixed-selective cortical populations. Furthermore, it is possible that regions more directly involved in feature-based attentional control, such as the superior colliculus, parvocellular layers of the lateral geniculate nucleus, ventral pulvinar, or the ventral prearcuate (VPA) region of prefrontal cortex (Schneider, 2011; Bichot et al., 2015; Lin et al., 2025), would show stronger RPE modulation.

Across the different metrics and monkeys we sometimes see different RPE functions, such as Absolute or Quadratic, performing similar to the Switch model. It is possible that the true RPE relationship is not any one of our pre-defined options but rather a more complex function. To address this, we attempted a complementary data-driven approach (not shown), using deep symbolic regression to derive the relationship between value, RPE, and attention by directly fitting monkey trialwise choices. However, this approach faced optimization challenges and did not find a better fit than the results presented here.

More broadly, this work contributes to theories about value-based attention, which argue that reward learning and attentional selection are not independent but dynamically interact to shape behavior (Leong et al., 2017; Anderson, 2016). Our results extend these theories by specifying the mathematical functions through which RPE and value can modulate covert feature-based attention. However, several limitations remain. First, our modeling and neural analyses are restricted to one task and dataset. It is worthwhile to test if our results hold in other learning tasks. Second, the neuron analyses were limited to single neuron correlations in a set of frontal and parietal brain regions. Datasets with larger population level simultaneous recordings, especially from visual areas such as V4 during learning, may help reveal more about the driving factors for attention. Finally, while our model captures learning dynamics at the behavioral level, it abstracts away from circuit-level mechanisms that may implement such attentional gain control. For example, neuromodulatory systems (e.g., dopamine, noradrenaline) are known to be involved in attention and reward prediction error (Avery and Krichmar, 2017; Diederen and Fletcher, 2021). Future work will need to use computational modeling approaches to gain insights into the circuit-level mechanisms implementing these RPE-regulated attention-learning interactions.

## 5 Extended Data

**Figure 8:**
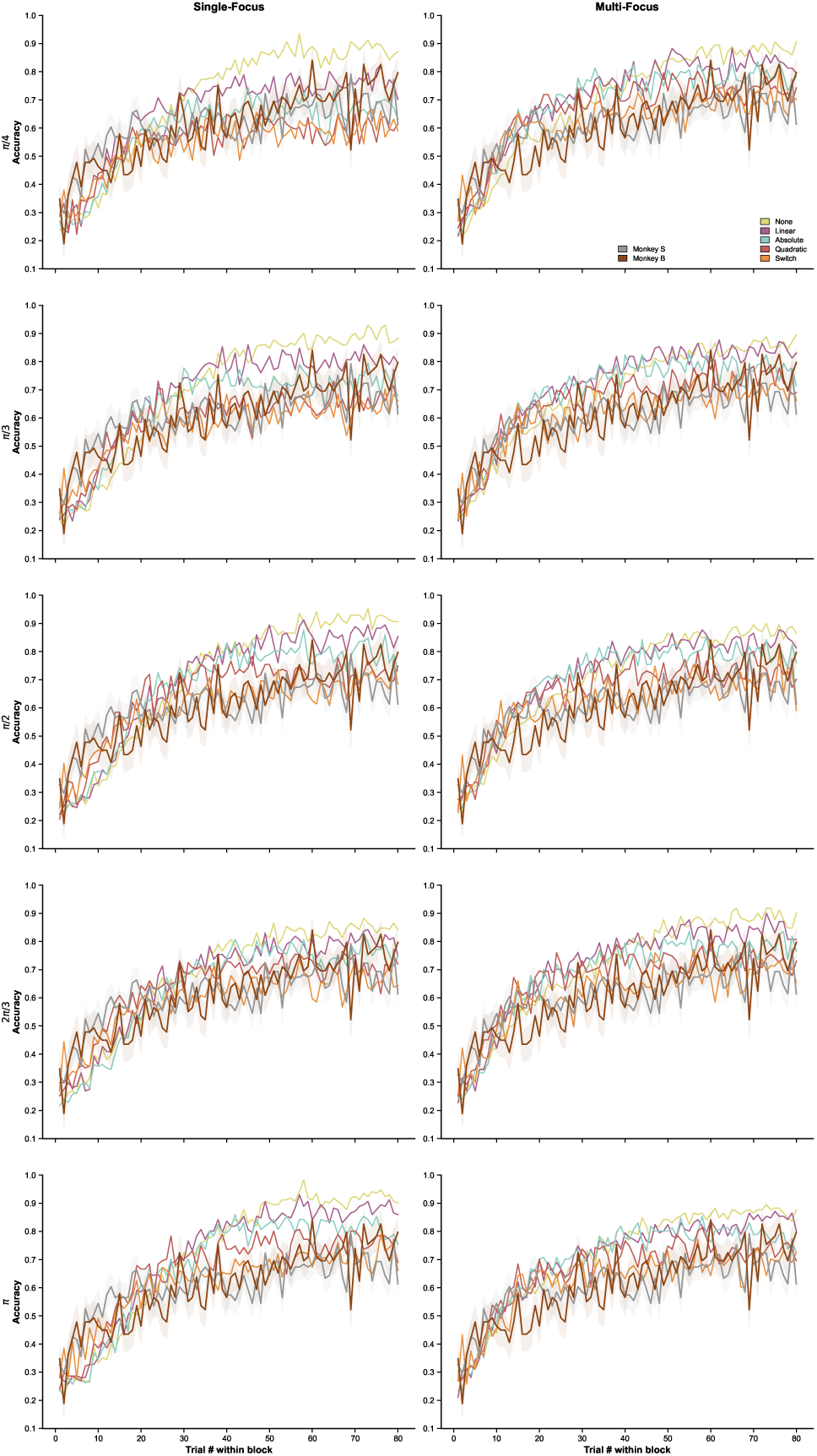
Effect of attention width on model learning curves. Figure shows block-aligned accuracy (mean ± SEM across 20 simulation seeds) as a function of trial number within a block, for five RPE-modulated attention models (None, Linear, Absolute, Quadratic, Switch). Columns separate single-focus (left) and multi-focus (right) architectures. Empirical learning curves for Monkey B (brown; 69 blocks) and Monkey S (gray; 101 blocks) are overlaid for comparison, with SEM computed across blocks. Attention width controls the angular breadth of the attentional gain field applied to feature-space representations, with narrower widths producing more selective enhancement of attended features. Across widths (*π/*4*, π/*3*, π/*2, 2*π/*3*, π*), model learning curves remain qualitatively similar to empirical data, indicating that the behavioral predictions of RPE-modulated selective attention are robust to variation in this parameter.

**Figure 9:**
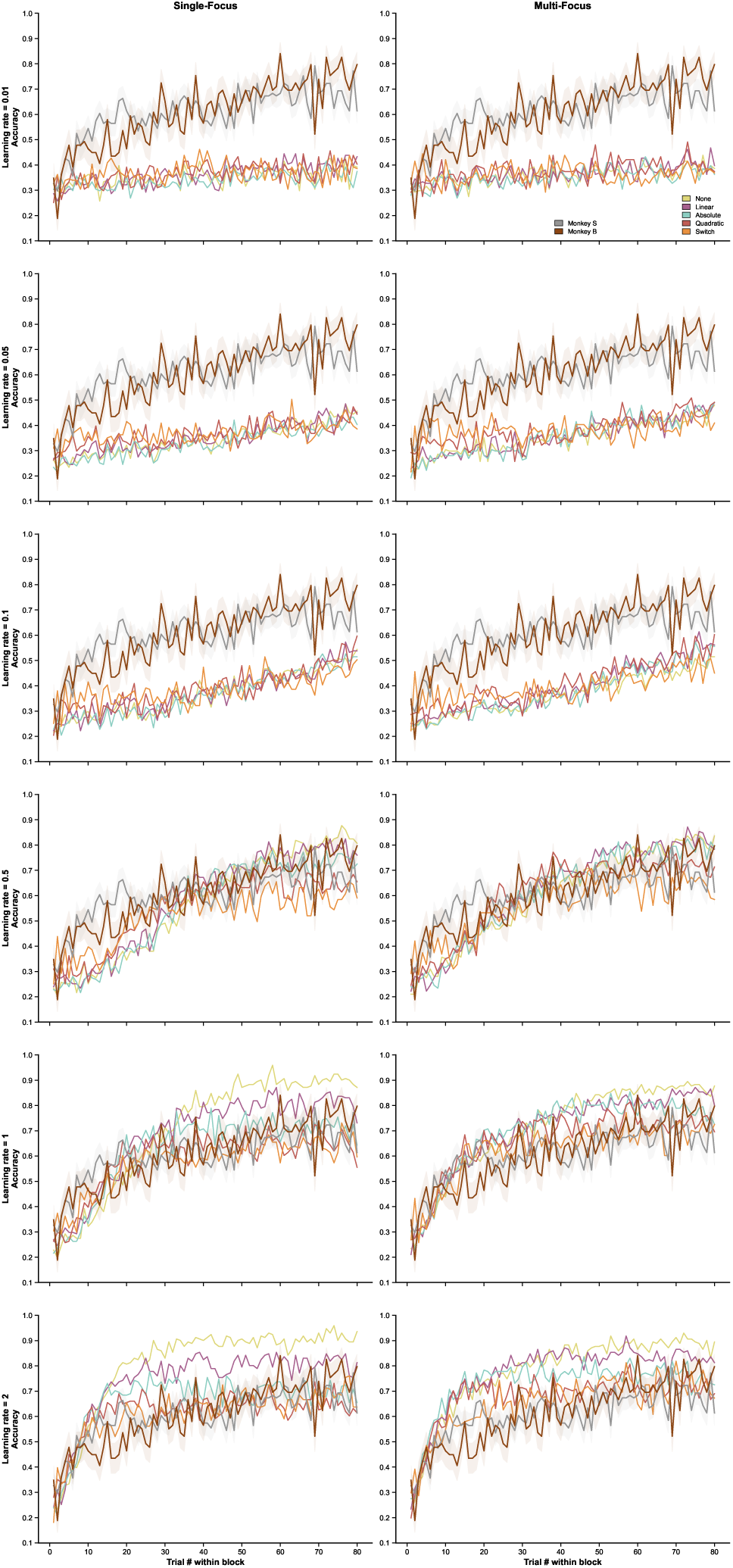
Effect of learning rate on model learning curves. Each panel shows block-aligned accuracy (mean ± SEM across 20 simulation seeds) as a function of trial number within a block, for five RPE-attention models (None, Linear, Absolute, Quadratic, Switch). Columns separate single-focus (left) and multi-focus (right) architectures. Empirical learning curves for Monkey B (brown; 69 blocks) and Monkey S (gray; 101 blocks) are overlaid for comparison, with SEM computed across blocks. The learning rate parameter (*α*) controls the step size of value updating 4.7, tested across a wide range (0.01 to 2). At very low learning rates (*α* = 0.01), models learn too slowly to perform above chance level (33%). Higher values (*α* = 0.5 to 2) produce model trajectories most consistent with the empirical data, confirming that the qualitative pattern of results reported in the main text is not an artifact of a narrowly tuned learning rate.

**Figure 10:**
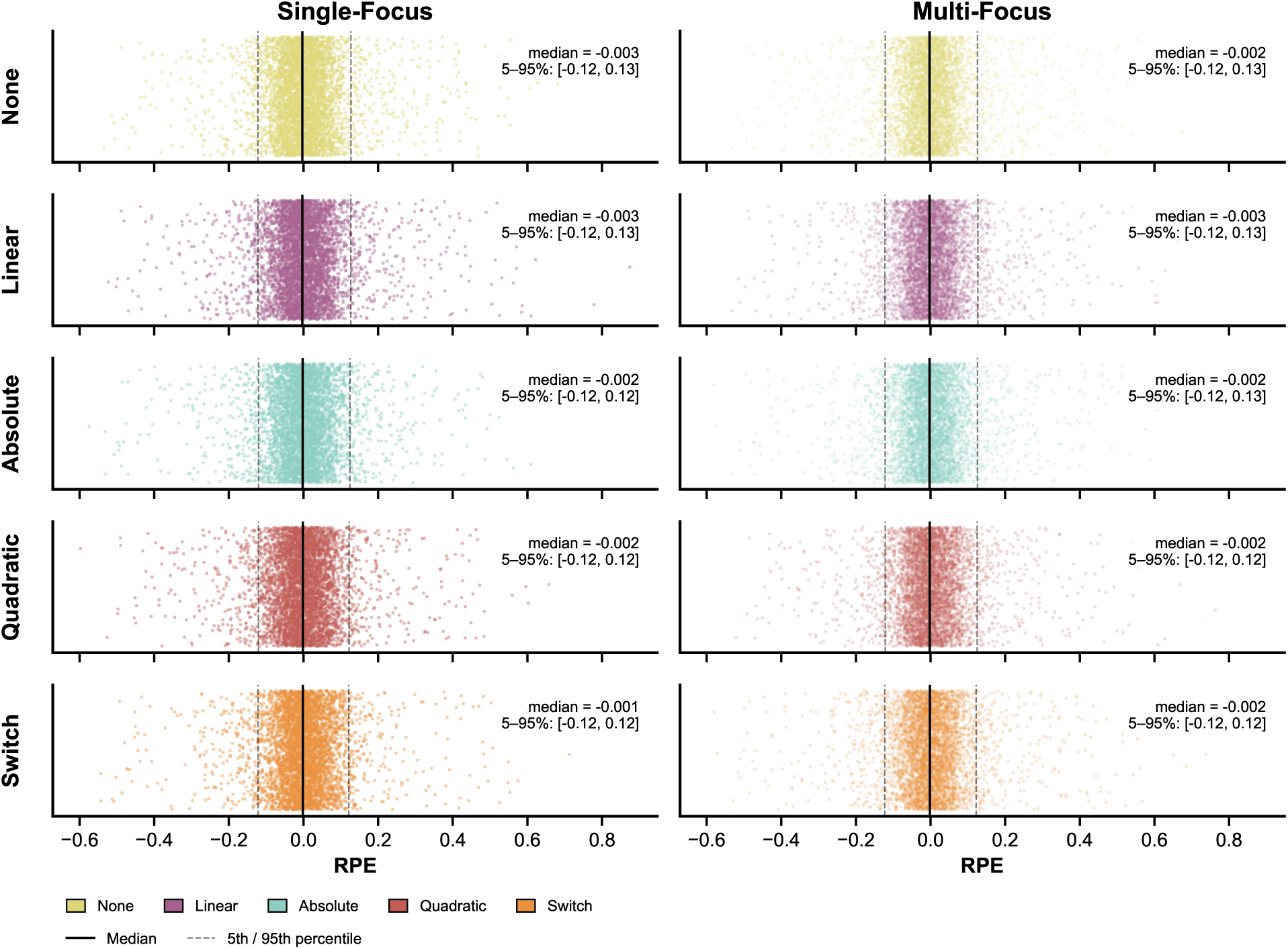
Empirical distribution of reward prediction error (RPE) values across model configurations. Each panel displays a scatter plot of RPE values for a given model, organized by RPE-attention mapping type (rows: None, Linear, Absolute, Quadratic, Switch) and attention architecture (columns: Single-Focus, Multi-Focus). Solid vertical lines indicate the median RPE; dashed vertical lines mark the 5th and 95th percentiles. Across all ten model configurations, RPE values occupied a narrow and consistent range (5–95th percentile: approximately [−0.12, 0.13]).

## Acknowledgments

This work was supported by New York University through the Mc-Cracken Fellowship (M.L.) and The Training Program in Computational Neuroscience (TPCN) under project number R90DA060339 (to M.L.). We thank the Buschman lab for openly providing their data and code, which were essential for this work. We also thank the members of the Lindsay lab for providing thoughtful advice and support for this project.

